# Psychosocial experiences are associated with human brain mitochondrial biology

**DOI:** 10.1101/2023.10.06.559575

**Authors:** Caroline Trumpff, Anna S Monzel, Carmen Sandi, Vilas Menon, Hans-Ulrich Klein, Masashi Fujita, Annie Lee, Vladislav Petyuk, Cheyenne Hurst, Duc M. Duong, Nicholas T. Seyfried, Aliza Wingo, Thomas Wingo, Yanling Wang, Madhav Thambisetty, Luigi Ferrucci, David A. Bennett, Philip L. De Jager, Martin Picard

## Abstract

Psychosocial experiences affect brain health and aging trajectories, but the molecular pathways underlying these associations remain unclear. Normal brain function relies heavily on energy transformation by mitochondria oxidative phosphorylation (OxPhos), and two main lines of evidence bi-directionally link mitochondria as both targets and drivers of psychosocial experiences. On the one hand, chronic stress exposure and possibly mood states alter multiple aspects of mitochondrial biology; and on the other hand, functional variations in mitochondrial OxPhos capacity alter social behavior, stress reactivity, and mood. However, knowledge on whether positive or negative psychosocial exposures and experiences are linked to mitochondrial biology in the human brain is currently unknown. By combining longitudinal antemortem assessments of psychosocial factors with postmortem brain (dorsolateral prefrontal cortex) proteomics in older adults, we found that positive experiences (e.g. higher well-being) are linked to greater abundance of the mitochondrial OxPhos machinery, whereas negative experiences (e.g. higher negative mood) are linked to lower OxPhos protein content. Combined, psychosocial factors explained 18% of the variance in the abundance of OxPhos complex I, the primary biochemical entry point that energizes brain mitochondria. To increase the sensitivity of our approach, we next interrogated mitochondrial psychobiological associations in specific neuronal and non-neuronal brain cells with single-nucleus RNA sequencing. These results revealed strong cell type specific associations, particularly between positive psychosocial experiences and molecular mitochondrial phenotypes in glial cells, whereas neurons tended to show opposite associations. Accordingly, in bulk transcriptomic analyses where all cells are pooled, these RNA-based associations were masked. Thus, our results highlight the likely underestimation of effect sizes in bulk brain tissues, and document novel cell type specific mitochondrial psychobiological associations in the human brain. Cell type specific mitochondrial recalibrations represent a potential psychobiological pathway linking positive and negative psychosocial experiences to human brain biology.

**Significance statement:** Psychosocial experiences predict health trajectories, but the underlying mechanism remains unclear. We found that positive psychosocial experiences are linked to greater abundance of the mitochondrial energy transformation machinery, whereas negative experiences are linked to lower abundance. Overall, we found that psychosocial experiences explain 18% of the variance in abundance of complex I proteins, the main entry point of the mitochondrial oxidative phosphorylation (OxPhos) system. At single-cell resolution using single nucleus transcriptomics, positive psychosocial experiences were particularly related to glial cell mitochondrial phenotypes. Opposite associations between glial cells and neurons were naturally masked in bulk transcriptomic analyses. Our results suggest that mitochondrial recalibrations in specific brain cell types may represent a potential psychobiological pathway linking psychosocial experiences to human brain health.

## Introduction

Psychosocial experiences predict health outcomes and longevity through mechanisms that remain unclear, but likely involving the brain (1). Positive psychosocial factors such as larger social network or greater purpose in life are associated with reduced risk of mortality and higher cognitive functioning at an older age (e.g. (2–6)), while negative factors, such as social isolation, loneliness, and depression are associated with increased risk for cognitive impairments (e.g. (7–9)). However, the sub-cellular and molecular pathways that transduce psychosocial exposures and affective states into lifespan-altering cellular processes within the brain and body remain unclear.

On the one hand, mitochondria are targets of stress and represent a potential pathway for the biological embedding of psychosocial exposures (10–14). The brain sustains its function by relying on energy (i.e., ATP) produced through oxidative phosphorylation (OxPhos) within mitochondria (15–19). Stressful experiences and chronic psychosocial stressors are believed to trigger neuroendocrine, molecular, and functional recalibrations that affect macroscopic brain properties (leading, for example, to hippocampal atrophy (20) and cortical thinning (21)), as well as sub-cellular and organellar structures, including mitochondria, that support numerous neuronal and glial functions (11, 22). Preclinical studies have shown that acute and chronic psychological stress affect mitochondrial content and functions across multiple tissues (e.g. (23–25); for a review see (10)). In bulk analyses of rodent brains, chronic stress affects the expression of mitochondrial gene expression involved in energy transformation (26, 27) as well as the abundance of proteins involved in mitochondrial energy metabolism, oxidative stress, and mitochondrial import and transport (27–29), likely signaling increased energy demand. In humans, chronic psychological stress and psychopathology are associated with molecular and functional alterations in specific aspects of mitochondrial biology measured in blood cells, including whole blood mtDNA copy number (mtDNAcn) (30, 31), enzymatic activities (32, 33), and cellular respiration (34–38), suggesting that psychosocial experiences affect mitochondrial biology across multiple tissues. These data implicate energetic and mitochondrial recalibrations as features of cellular responses to mental stress and possibly other psychosocial exposures.

On the other hand, mitochondrial biology also may drive behavior and affective experiences. In preclinical rodent models, mitochondrial DNA defects alter physiological stress-reactivity (39–41) and influence mood-like behaviors (40, 42). Alteration in mitochondrial OxPhos in defined brain regions (nucleus accumbens, prefrontal cortex) influences social behavior as well as anxiety-like and depression-like behaviors ((27, 43–48), reviewed in (49)). In syngenic animals (with the same genome), we also found that differences in mitochondrial energy production capacity in brain tissue accounted for a large fraction (20-45%) of animal-to-animal differences on anxiety-like behaviors and social interaction/avoidance (50). In humans, psychopathology, autism, and/or neurodegeneration are also related to alterations in brain mitochondrial OxPhos components and mtDNA (e.g. (27, 51) see (52) for a review). Thus, while chronic stress can affect brain mitochondrial biology, mitochondrial biology may also directly influence mood and behavior.

Existing human studies documenting a connection between daily psychosocial experiences and mitochondria have mostly been conducted in peripheral immune cells (32, 34–37). While these and other studies identified mitochondrial biology as a potential psychobiological pathway, findings in leukocytes are limited by high inter-individual variation in cell type abundances, rapid cell turnover of mitotic cell (sub)populations, and dynamic immune processes that influence mitochondrial biology but say little about the “health” or systemic properties of mitochondria themselves (53). In comparison, the brain contains neurons and glial cells that exist in relatively stable proportions (compared to immune cells) across the lifespan. Moreover, mitochondrial metabolism specifically directs cellular activities such as neuronal excitability and presynaptic neurotransmitter release (54), neurogenesis (55), as well as glial cell properties including astrocyte proliferation (56), microglia activation (57, 58), and oligodendrocyte differentiation (59). For reasons that remain unknown but may relate to the preferential oxidation of glucose and unique metabolic requirements for OxPhos complex I in brain tissue, the brain is particularly vulnerable to OxPhos complex I defects in human genetic diseases and animal models (60, 61). Neuronal developmental rates and cellular aging trajectories also are under mitochondrial regulation (62, 63), opening the possibility that mitochondrial alterations could influence brain aging and the risk of neurodegenerative diseases (64).

Given the central role of the brain in the *processing* of psychosocial exposures as well as in the *production* of affective experiences, it represents the most relevant tissue to investigate evidence of mitochondria psychobiological associations. To address this question, we examine how prospectively-collected positive and negative psychosocial factors relate to post-mortem protein and transcript abundance of known mitochondrial pathways in hundreds of human brains (dorsolateral prefrontal cortex) (Fig. 1A), in bulk brain tissue and in single cells.

**Fig 1.**
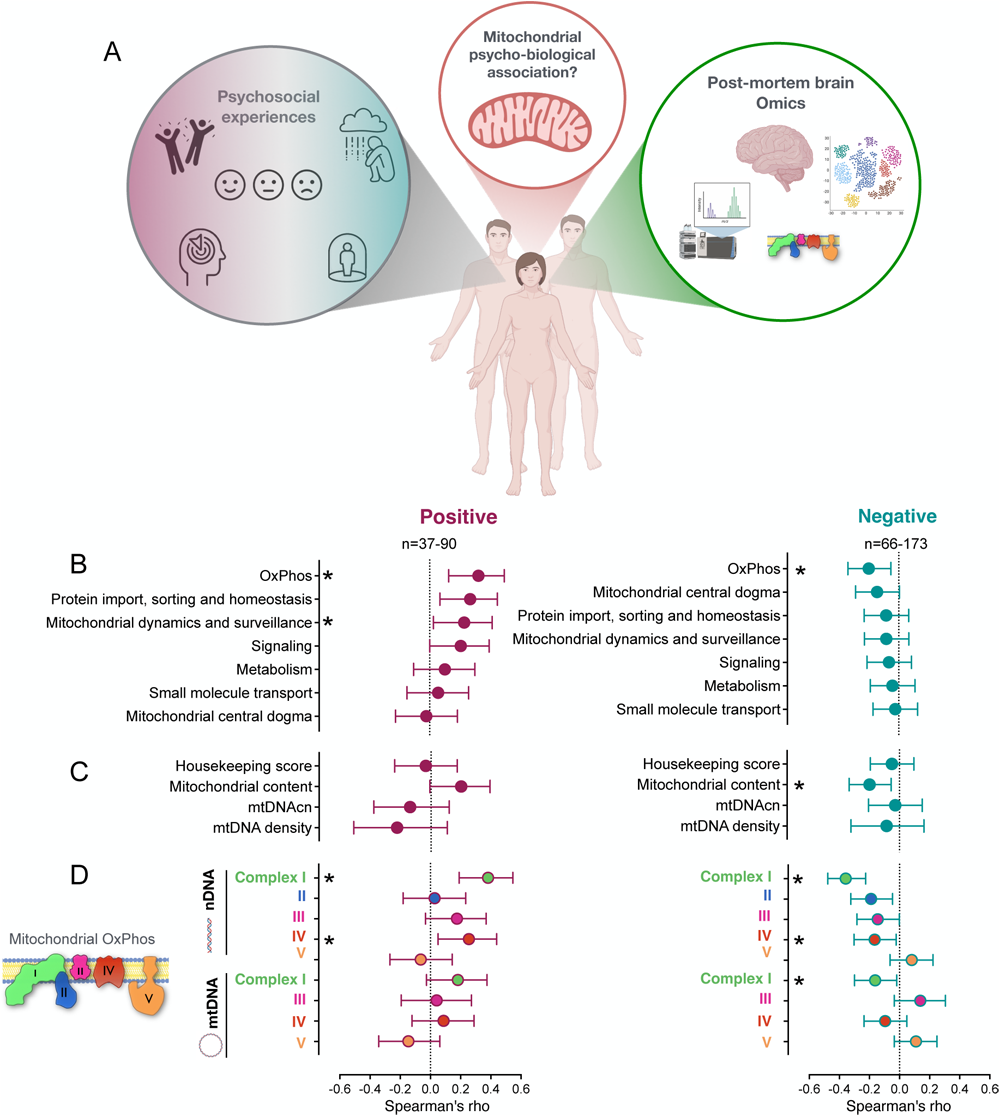
Psychosocial experiences and the mitochondrial brain proteome. (**A**) Study design: the ROS and MAP cohorts include psychosocial data collected prospectively as well as post-mortem brain multi-omics including TMT Proteomics (for more details see Supplemental figure S1A). Study goal: we assessed the relationship between positive and negative psychosocial experiences and post-mortem dorsolateral prefrontal cortex (DLPFC) mitochondrial biology. (**B**) Effect size (spearman’s rho (95% CI)) for the association between positive and negative psychosocial scores and average protein abundance for the 7 main mitochondrial pathways referenced in MitoCarta 3.0. (level 2 and 3 sub-categories are shown in Supplemental Fig. S6-7, for study gene coverage see Supplemental table S2); (**C**) cellular housekeeping score, mitochondrial content score (average of all available proteins referenced in MitoCarta 3.0.), mitochondrial DNA copy number (mtDNAcn) and mtDNA density (mtDNAcn/mitochondrial content score); (**D**) and mitochondrial OxPhos summary scores adjusted for mitochondrial content. Detailed results are shown in Supplemental table S3A. Protein abundances were regressed for age at death, postmortem interval, study and batch prior to analysis.

## Results

### Psychosocial experiences are associated with OxPhos protein abundance

We used data from the ROS and MAP cohort studies (65, 66) (study design and cohort characteristics are shown in Fig 1A,B, Fig. S1 and Table S1). This dataset includes longitudinal prospectively acquired psychosocial data, complemented by untargeted quantitative proteomics offering a broad coverage of mitochondrial proteins from post-mortem brain dorsolateral prefrontal cortex (DLPFC), a brain area involved in executive functions and emotional regulation (67), and sensitive to psychological stress exposure (27, 68–70).

We approached mitochondria as multifaceted organelles that are best studied as multiple functional domains (32, 71, 72), rather than by a single indicator. To capture interpretable biological signatures of mitochondrial health, we aggregated >1,100 mitochondrial genes as *MitoPathways* (73) (for study coverage see Table S2). Because each mitochondrial pathway reflects the average of several functionally related genes, *MitoPathways* are both more biologically *interpretable* and also more statistically *robust* than single-gene or single-protein approaches.

We used a similar approach to integrate psychosocial experiences, defined here as the psychosocial exposures and psychological states in an individual’s outer and inner world. This includes *positive factors:* social network size, late life social activity, purpose in life, time horizon, well-being; and *negative factors:* adverse childhood experience, negative life events, social isolation, depressive symptoms, negative mood, perceived stress. From these factors, we built a positive and a negative score within which longitudinal variables were averaged across the follow-up period (average duration: 4.3 years ± 2.2 (S.D.), range 1-10 years; see Fig. S2 for score composition and inter-correlations), providing more stable estimates of each person’s psychosocial and mental states prior to death, which can then be related to mitochondrial biology.

Mitochondrial proteomics and psychosocial data were available for 400 participants (281 women, 119 men). To limit the dimensionality and complexity of our analyses, we pre-selected 7 main *MitoPathways* on the basis of previous literature and mechanistic upstream drivers of OxPhos function: “OxPhos”, “protein import, sorting and homeostasis”, “mitochondrial dynamics and surveillance”, “signaling”, “metabolism”, “small molecule transport” and “mitochondrial central dogma” (73). Correlation analyses of the protein-based MitoPathway scores with either positive or negative psychosocial scores showed that OxPhos consistently exhibited the strongest psychobiological associations (Fig. 1B).

Positive psychosocial factors were positively associated with “protein import, sorting and homeostasis” (rho=0.27; 95%CI=0.07 to 0.45), “mitochondrial dynamics and surveillance” (rho=0.23; 95%CI=0.03 to 0.42), and OxPhos protein abundance (rho=0.33; 95%CI=0.13 to 0.50). On the other hand, negative psychosocial score was negatively associated only with OxPhos (rho=-0.27; 95%CI=-0.40 to -0.14), indicating that individuals reporting more negative experiences have fewer mitochondrial OxPhos proteins in the DLPFC.

In stratified analyses, the associations with OxPhos were found across both men (positive: rho=0.45; 95%CI=0.10 to 0.70, negative: rho= -0.17; 95%CI=0-.41 to -0.17) and women (positive: rho=0.28; 95%CI=0.03 to 0.50, negative: rho=-0.30; 95%CI=-0.45 to -0.13). Similar psychosocial-OxPhos were observed for individuals with cognitive impairment (positive: rho=0.47; 95%CI=0.02 to 0.76, negative: rho=-0.30; 95%CI=-0.58 to 0.04) or without cognitive impairment (positive: rho=0.34; 95%CI=0.05 to 0.57, negative: rho=-0.39; 95%CI=-0.54 to -0.21) (Fig. S2, Table S3C), supporting the generalizability of these findings linking positive and negative to more and less DLPFC OxPhos protein content, respectively.

### Psychobiological OxPhos associations in BLSA and with TMT proteomics

In a much smaller study cohort with multiple brain areas, the Baltimore Longitudinal Study of Aging (BLSA) (n=47, 36% female, see Fig. S3 and Table S2 for cohort characteristics), there were no significant associations between psychosocial factors and mitochondrial pathways in the DLPFC. However, in the precuneus area of the same participants, positive psychosocial experiences were positively associated with mitochondrial dynamics and surveillance protein abundance (rho=0.47; 95%CI=0.14 to 0.71; n=15), while the reverse was found for negative psychosocial experiences (rho=-0.34; 95%CI=-0.56 to -0.07; n=26) (Fig. S4). Thus, while not a straightforward validation of our findings in the ROSMAP cohort, the directionality of findings aligns for related but different *MitoPathways*. Together with other work showing mitochondrial variations across mouse brain areas (50), this calls for future studies assessing mitochondrial psychobiological associations across different brain regions.

In ROSMAP, we found no significant relationship between psychosocial scores and DLPFC mitochondrial DNA copy number (mtDNAcn), mtDNA density (mtDNAcn/mitochondrial content score), or a general mitochondrial housekeeping score (Fig. 1C). Although people who reported more positive and negative experiences tended to have greater (rho=0.20; 95%CI=-0.01 to 0.39) and lower (rho=-0.33; 95%CI=-0.33 to -0.05) total estimated mitochondrial mass, respectively (Fig. 1C), correcting OxPhos abundance for mitochondrial mass showed that the psychosocial experiences and OxPhos associations were not fully explained by differences in total mitochondrial content (Fig. 1D). These results thus largely rule out that psychosocial experiences would be only associated with unspecific changes in mitochondrial mass, for example driven by a general increase or decrease in energy demand. Instead, our results, taking into account both OxPhos proteins and mitochondrial content, suggest that psychosocial experiences are linked to differences in OxPhos proteins on a per-mitochondrion basis (i.e., a qualitative change in mitochondrial health).

To examine this possibility with the highest possible degree of specificity, we examined more specific *MitoPathways* quantifying specific components of the OxPhos system, including complexes I through V. The strongest association with psychosocial exposures was for nDNA-encoded subunits belonging to complex I (positive: rho=0.38; 95%CI=0.19 to 0.55, negative: rho=-0.36; 95%CI=-0.48 to -0.23) (Fig. 1D, Fig. S5-7). OxPhos complex I is the major entry point for electrons into the electron transport chain (ETC), and is the OxPhos component whose abundance and impaired activity in the prefrontal cortex has been the most frequently associated with neurodegeneration and neurodevelopmental disorders (51, 52).

We then sought to confirm these data in a larger selected reaction monitoring proteomics (SRM, n=1,208) ROSMAP dataset, which contains only a small subset of targeted mitochondrial proteins (16 OxPhos proteins total, vs 128 for the TMT dataset). The associations between psychosocial factors and complex I nDNA-encoded subunits showed the same direction, but the effects were only significant for negative factors (rho=-0.24; 95%CI=-0.13 to -0.03) (Fig. S8). This suggests two things: first, that our findings partially generalize in a larger, heterogenous sample; and second, in keeping with the multifaceted nature of mitochondria and multi-protein nature of OxPhos complexes (50), that the mitochondrial indices built from comprehensive sets of proteins (in the TMT DLPFC dataset, Fig. 1) covering >90% of OxPhos proteins might be required to sensitively detect mitochondrial psychobiological associations.

### Domain-specific OxPhos psychobiological associations

Next, we dissected the separate contributions of each instrument that make up the positive and negative psychosocial scores to see which contributed the most to the psychobiological associations with OxPhos protein abundance. For the positive psychosocial score, the significant measure contributing the most the OxPhos association were well-being (rho=0.31; 95%CI=0.11 to 0.48), which was assessed according to six major themes (purpose in life, personal growth, positive relations with others, self-acceptance, autonomy environmental mastery) (74, 75), and late-life social activity (rho=0.20; 95%CI=0.06 to 0.32), which captures how often participants engaged in activities involving social interaction (76). For the negative score, negative mood (rho=-0.24; 95%CI=-0.39 to -0.08) and negative life events (rho=-0.19; 95%CI=-0.34 to -0.03) had the strongest effect sizes (Fig. S9, stratified analysis are shown in Fig. S10, Table S3G). Thus, both individual experiences (well-being and mood) and objectifiable factors (social activity and life events) relate to DLPFC brain mitochondrial biology.

To quantify the relative and combined contributions of psychosocial experiences on mitochondrial proteins, we performed multiple linear regressions controlling for relevant covariates. Positive and negative psychosocial factors explained 11% (p=0.0015) and 12% (p<0.0001) of the variance in nDNA-encoded subunits of complex I abundance, respectively (Fig. S11). Since individuals with more positive experiences may also report less negative experiences, we created a combined score taking into account their shared variance. The psychosocial factors assessed in this study explained 18% of the variance in nDNA-encoded complex I subunit abundance (r^2^=0.18, p<0.0001) (Fig. 2), indicating a large potential cross-talk between psychosocial experiences and brain mitochondrial OxPhos capacity.

**Fig 2.**
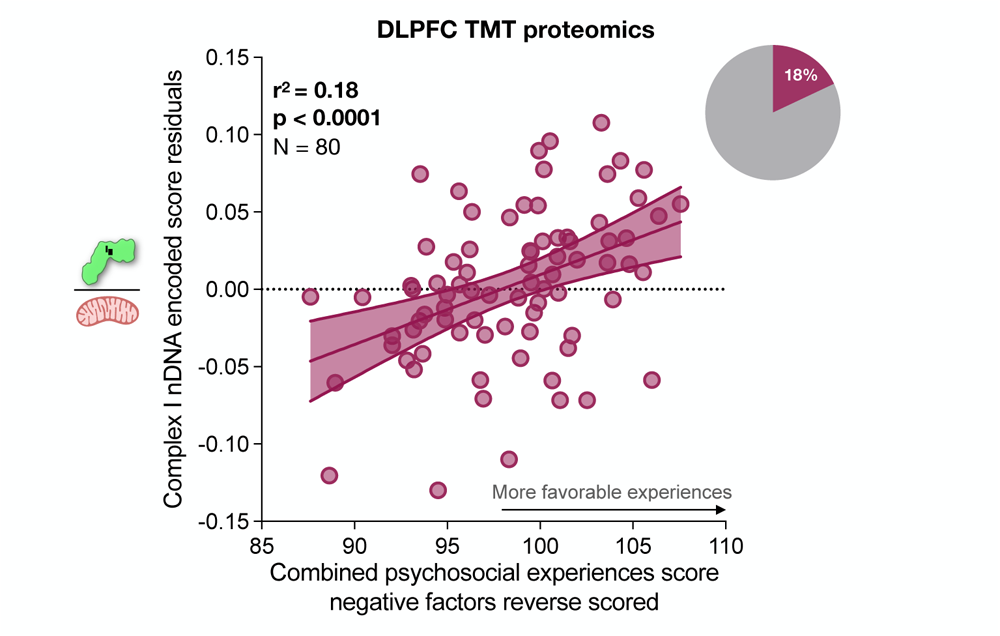
Positive psychosocial experiences account for a substantial proportion of inter-individual differences in human DLPFC mitochondrial complex I protein abundance. The total proportion of variance in complex I nDNA-encoded protein abundance (adjusted for mitochondrial content) attributable to psychosocial experiences was computed using positive and negative (reversed) variable scores (excluding social network size). Results from multivariate linear regressions adjusting for sex and cognitive status; protein abundances were regressed for age at death, postmortem interval, study (ROS and MAP), and technical batches.

### Mitochondrial psychobiological associations are cell type specific

To examine if the proteomic findings above (reflecting the OxPhos machinery) were also observed at the level of gene expression (largely reflecting the active cellular programs rather than the machinery), we repeated the same analytical approach with bulk RNA-seq data. We analyzed three different brain areas: DLPFC (N=1,092), posterior cingulate cortex (PCC) (N=661), and anterior caudate (AC) (N=731) from the ROSMAP cohort. Unlike with proteomics, there were no consistent associations between positive nor negative experiences and RNA transcript-based mitochondrial pathways (Fig. S12-13). This suggested that either the mitochondrial psychobiological associations observed in the tissue proteome are limited to post-transcriptional factors (e.g., protein synthesis, stability, or other factors), or that the bulk transcriptome data is too noisy and does not provide the level of sensitivity or specificity required to detect true associations between psychosocial experiences and mitochondrial features.

Knowing that different cell types contain vastly different mitochondrial phenotypes (50, 53, 77), we therefore quantified mitochondrial phenotypes in specific brain cell types using single-nucleus RNA sequencing of ROSMAP DLPFC samples (n=424). Of the 87 distinct cell types and subtypes previously identified (78), we first examined mitochondrial psychobiological associations with transcript levels in cell types where the same *MitoPathways* as above were sufficiently well represented (>50% of participants and >50% coverage of mitochondrial genes). Based on these criteria, we included the following six major cell types: excitatory and inhibitory neurons, microglia, astrocytes, oligodendrocytes and oligodendrocyte progenitor cells (OPCs) (Fig. 3A).

**Fig 3.**
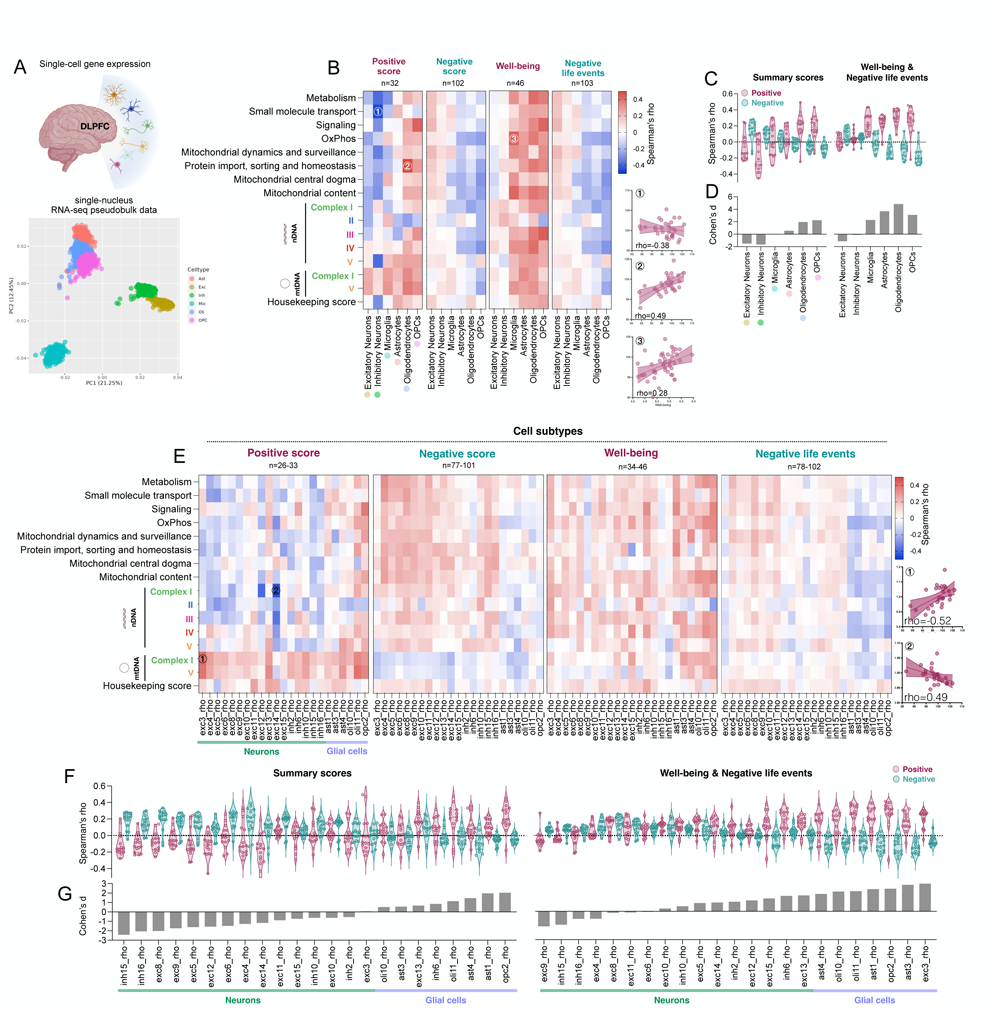
Psychosocial experiences and the mitochondrial brain transcriptome in specific cell types. (**A**) Study design: we assessed the relationship between positive and negative psychosocial experiences and post-mortem dorsolateral prefrontal cortex (DLPFC) mitochondrial transcript abundance in specific cell types (single-nucleus RNA-seq pseudobulk data) in the ROSMAP cohort. Effect size (spearman’s rho) for the association between positive and negative psychosocial scores and the 7 main MitoCarta 3.0. pathways scores, the cellular housekeeping score, the mitochondrial content score and the mitochondrial OxPhos summary scores displayed using (**B**) heatmaps and (**C**) violin plots (each dot represent one effect size, e.g. the association between positive experiences and metabolism). (**D**) Magnitude of difference (cohen’s d) between the average rho found for the association with positive vs negative psychosocial scores. (**E**), Same as in (B) for specific cell subtypes. (**F, G**) Same as in (C, D) sorted by effect size. Detailed results are shown in Supplemental table S4. Transcript abundances were regressed for age at death, postmortem interval, study, cognitive status and sex prior to analysis.

At this level of cellular resolution, we observed remarkable divergence in the magnitude and direction of the correlations between positive and negative experiences and *MitoPathways* DLPFC gene expression (Fig. 3B, Table S4B). Averaging all *MitoPathways* together to decrease false discovery rate, we found that individuals reporting more positive experiences had higher mitochondrial gene expression in both oligodendrocytes (rho^avg^=0.19) and OPCs (rho^avg^ =0.18). In contrast, positive experiences were negatively correlated with neuronal *MitoPathways*, both in excitatory (rho^avg^=-0.07) and inhibitory neurons (rho^avg^=-0.16).

These divergent neuron vs glial cell patterns were particularly apparent when examining the well-being scale alone. From the lowest to highest associations, we find neurons rho^avg^ =0.02 (range=-0.10 to 0.21), astrocytes rho^avg^ =0.23 (-0.03-0.34), microglia rho^avg^ =0.24 (0.06-0.43), oligodendrocytes and OPCs rho^avg^=0.28 (-0.22-0.45). Figure 3C presents the contrasts in standardized effect sizes between positive and negative experiences and all *MitoPathways*, for each major cell type. For this analysis, the null hypothesis is that both positive and negative exposures are not associated with *MitoPathway* expression (i.e., r=0.00) and that positive and negative scores or scales yield similar random distributions (i.e., effect size between positive and negative experiences is null (Cohen’s d=0). In contrast, glial cells exhibited very large contrasting effect sizes, indicating that psychosocial experiences are consistently associated in the same direction with multiple DLPFC single-cell *MitoPathways,* particularly among oligodendrocytes (Cohen’s d=1.95) and OPCs (d=2.25), but not in neurons (d<1.13).

We then expanded these analyses of mitochondrial psychobiological associations at the highest possible level of resolution across 23 cell subtypes having sufficient coverage (>50% of participants, >50% coverage of mitochondrial genes). Although underpowered for these analyses, as for our results among the six major cell types, these data also highlighted how individuals who report higher positive experiences tended to have greater *MitoPathway* gene expression in glial cells, but not in neurons (Fig. 3 E-G, Table S4C). Comparing nuclear vs mtDNA-encoded gene associations also suggested that mtDNA-encoded OxPhos subunits showed a positive association with psychosocial experiences across cell types (positive score: rho^avg^ = 0.23), while this was not the case for nDNA-encoded OxPhos subunits (positive score: rho^avg^ = -0.03) (Fig. 3 E), possibly reflecting differential regulation of nuclear and mitochondrial genomes.

Together, these single cell-level data provide two main insights. First, they show that mitochondrial psychobiological associations in the mitochondrial proteome may represent gross under-estimations of the true effect sizes. And second, that psychosocial factors have differential effects on mitochondrial biology in specific brain cells, or possibly, that mitochondria in glial cells contribute to affective experiences more than mitochondria in neurons.

## Discussion

In this study, we used longitudinal psychosocial data and post-mortem DLPFC proteomics in a sample of older adults to evaluate the association between psychosocial experiences and brain mitochondrial biology. Individuals reporting more positive experiences, such as greater well-being, had greater abundance of OxPhos complex I proteins, while the opposite effect was found for negative experiences. Considering their independent contributions (people feeling more positive may report fewer negative experiences) we find that ∼18% of the variance in complex I abundance between individuals was attributable to self-reported psychosocial experiences. This fraction of explained variance is substantial given the measurement errors inherent to self-reported questionnaires as well as to post-mortem brain omic measures. Moreover, extending these results using single nucleus transcriptomics results showed that the connection between psychosocial factors and mitochondrial biology may be either predominantly, or perhaps restricted to, non-neuronal glial cells. This suggests that the proportion of explainable variance in complex I proteins noted above is likely an underestimate, since it is derived from bulk brain tissue. Combined, our direct measurements of brain proteins (the active agents supporting electron transport chain function) together with our single-cell resolution transcriptomics data (showing large cell type specific associations) provide evidence linking psychosocial experiences and mitochondrial biology in the aging human brain.

Our findings linking psychosocial experiences to brain mitochondria agree with a vast body of animal literature documenting an effect of chronic stress exposure on various aspects of mitochondrial OxPhos. In animals, chronic stress leads to reduced protein expression specifically of complex I (79), complex I enzymatic activity (25, 80–83), and complex I-driven respiration (24). Thus, in line with our human brain findings, chronic stress may preferentially affect brain mitochondrial complex I biology for reasons that remain unclear.

Although animal studies document a causal effect of stress exposure on mitochondrial biology, our findings could reflect the reverse association. DLPFC mitochondrial bioenergetics could contribute to shaping affective experiences, such as one’s state of well-being reported on questionnaires. This interpretation is supported by a growing body of work in rodents and humans indicating that mitochondrial OxPhos defects can alter behavior (e.g. (24, 45, 48)). Social dominance, depressive, and anxiety-related behaviors have been associated with variation in mitochondrial OxPhos capacity (43–46, 50). In the human brain, post-mortem studies show reduced complex I activity in relation to depression (see (52) for a review), and individuals with mitochondrial DNA defects leading to reduced OxPhos capacity exhibit a higher incidence of depressive symptoms compared to the general population (84). Using non-invasive proton magnetic resonance spectroscopy imaging in humans, a study also found that trait anxiety is associated with downregulated nucleus accumbens taurine levels (85), a metabolite essential for complex I activity (86, 87). Finally, complex I occupancy quantified by positron emission tomography (i.e., a proxy for protein abundance) was lower in the anterior circulate cortex of adults with autism spectrum disorder than controls, and lower abundance was correlated with more severe social communication deficits (51), thus suggesting that variation in mitochondrial energy production capacity in specific brain regions may influence social behaviors.

The brain, and in particular the DLPFC area, is directly relevant to the embedding of psychosocial experiences as it is involved in executive functions and emotional regulation. Sustained cognitive demand directly affects brain DLPFC metabolism characterized by glutamate accumulation (88), which can have neurotoxic effects when chronically occurring such as when being exposed to a chronically stressful condition. Adversity and chronic stress (68), which overlap with the negative psychosocial constructs assessed in our study, lead to both structural and functional DLPFC changes such as reduction of grey matter volume (69) and/or impairment of brain functional connectivity involved in attention shifts (70). Our findings open the exciting possibility that alterations of mitochondrial OxPhos function could mediate the effects of social factors on brain energetics, and therefore affect downstream brain functions.

In relation to mitochondria in other tissues, our findings align and contrast with the mixed literature in immune cells. Consistent with the current brain study, we previously found that positive mood predicted higher OxPhos enzymatic activities in PBMCs on subsequent days (12 to 48 hours), but mitochondrial activities did not predict mood on subsequent days, suggesting mood-to-mitochondria directionality for these effects (89). Interestingly, the observed effect size in PBMCs was r^2^=0.13-0.16, comparable to r^2^=0.18 for the current results in the brain. Other studies have also found mitochondrial alterations to be a potential mechanism for the biological embedding of early adversity (90, 91). In contrast, a study found that childhood maltreatment was associated with higher mitochondrial respiration in live PBMCs (37), possibly as a result of increased pro-inflammatory activity, since activated cells producing high cytokine levels must consume more energy. Similarly, another study found that women experiencing greater stress quantified by allostatic load gave birth to children with greater mitochondrial content at age 3-6 yr (92), possibly reflecting the activation of energy-costly pro-inflammatory programs among immune cells (93, 94). Some of the divergences between these results in immune cells and our brain data could reflect the different cell types used, and/or more generally the nature of immune vs. brain cells.

An important caveat of our study is the unresolvable directionality of effects. We can conceive of four biologically plausible scenarios linking psychosocial and mitochondrial outcomes. First, it is plausible that psychosocial experiences affect brain activity and therefore directly shape mitochondrial biology. Second, one’s mitochondrial biology could affect behavior and perception of self-reported psychosocial experiences on questionnaires. Third, a bi-directional relationship may exist, such that chronic stress exposure directly affects an individual’s mitochondrial biology and subsequently affects their perception of social events. And fourth, other factors such as environmental toxicant could affect both mitochondria and psychosocial experiences through independent mechanisms. However, the emerging picture in the literature is that all those pathways are interactive and, thus, our results may reflect the outcome of those complex interactions.

While our study design is strengthened by integrated indices from prospectively acquired psychosocial data, as well as *post-mortem* proteomics and single nucleus gene expression data in a large sample of individuals, it also presents several technical limitations. Proteomics lacks sensitivity to detect proteins present at very low abundance (including some mitochondrial proteins) and highly hydrophobic mitochondrial proteins embedded within lipid membranes (95). Because the enzymatic activities of proteins are post-translationally regulated by several factors, protein abundance also may not directly translate into mitochondrial functions and behaviors (71). Future studies should complement static, omic-based measures of molecular mitochondrial features with direct measures of mitochondrial activities and functions. *Post-mortem* gene expression may also only poorly reflect gene expression in the living human brain, calling for studies of living samples where possible (96), although achieving the scale required to address psychobiological hypotheses may be challenging. Other limitations include the lack of psychosocial measures proximal to death in some participants with AD (not able to provide questionnaire-based data), which we expect to contribute noise to our analyses, likely resulting in underestimated psychobiological associations. Finally, the use of self-reported measures of psychosocial experience is not without limitations (e.g., high variability, desirability bias), but the prospective nature of the assessments in this study likely prevented recall bias.

In summary, the present study is the first to our knowledge to link individual psychosocial experiences (antemortem) with human brain mitochondrial biology (post-mortem). Our results document a robust psychobiological association between psychosocial experiences and brain DLPFC mitochondrial OxPhos complex I. Moreover, single-nucleus studies of psychosocial-mitochondrial associations suggest that glial and neuronal cell subtypes may either respond to or contribute to human psychosocial experiences in opposite ways. Cell type specificity therefore represents an important concept to integrate into our emerging understanding of the processes linking psychosocial factors, brain energetics, and mitochondria. While the directionality of these effects and the underlying mechanisms remain to be examined in detail, our results are consistent with the notion that recalibrations in mitochondrial energetics may transduce the effects of psychosocial exposures into molecular and biological changes that shape brain health and aging.

## Materials and Methods

### Study participants

We used data from ROSMAP (N=400) (65, 66, 97) and BLSA (N=47) (98–100), (99–101). These studies have measured post-mortem DLPFC protein abundance using untargeted proteomics. Clinical characteristics of the study participants included in the analysis are shown in Table S1.

### ROSMAP

We repurposed post-mortem brain proteomics and RNAseq data from two cohort studies of adults, the Rush Memory and Ageing Project (MAP) and the Religious Orders Study (ROS) (65, 102, 103). ROS study participants are catholic nuns, priests, and brothers, from about 40 groups across the United States. MAP study participants were enrolled primarily from about 40 retirement communities throughout northeastern Illinois, with additional participants enrolled through home visits. Participants were free of dementia at study enrolment and agreed to annual evaluations and brain donation on death. The ROS and MAP studies were approved by the Institutional Review Board of Rush University Medical Center. All participants signed an informed consent, Anatomical Gift Act, and a repository consent to share data and biospecimens.

Alzheimer’s Disease (AD) clinical diagnosis proximate to death was based on a review of selected annual clinical data after death by the study neurologist who did not have access to neuropathologic data. Cognitive status was defined as no cognitive impairment (NCI), mild cognitive impairment (MCI), or Alzheimer’s dementia (AD) from the final clinical diagnosis of dementia, as reported previously (104–106). Post-mortem AD pathology was measured as described previously (104, 107), and AD classification was defined based on the National Institutes of Ageing-Reagan criteria (108). The NIA-Reagan criteria for the pathologic diagnosis of AD (high or intermediate likelihood of AD) (104) and final clinical diagnosis of cognitive status (AD and no other cause of cognitive impairment) was used to define pathologic AD as described previously (109). MCI refers to participants with cognitive impairment but who did not meet criteria for dementia.

### BLSA

The National Institute on Aging’s Baltimore Longitudinal Study of Aging (BLSA) was used as an additional study cohort. TMT proteomic data from DLPFC brain tissue was available for 13 cognitively healthy individuals and 34 AD cases (36% women). The BLSA study was approved by the Institutional Review Board and the National Institute on Aging. Human research at the National Institutes of Health (NIH) and the BLSA participants provided written informed consent (98).

Post-mortem neuropathological evaluation was conducted at the Johns Hopkins Alzheimer’s Disease Research Center with the Uniform Data Set. Amyloid plaque distribution was assessed according to the CERAD criteria and neurofibrillary tangle pathology was assessed with the Braak staging. Participants were classified as cognitively healthy if they were evaluated as cognitively healthy within 9 months of their death and presented low CERAD (0.13 ±0.35) and Braak (2.26 ±0.94) measures of amyloid and tau neuropathology. Participants were categorized as AD cases when classified as demented at the last clinical research assessment, and when the brains showed high CERAD (2.9 ±0.31) and Braak (5.4 ±0.82) scores (consistent with moderate to severe neuropathological burden). Asymptomatic AD cases were participants evaluated as cognitively normal proximate to death, with brains presenting high CERAD (2.1 ±0.52) and moderate Braak (3.6 ±0.99).

### Psychosocial variables

#### ROSMAP

Psychosocial variables were either collected once or annually. Positive psychosocial exposures measured annually included social network size (110), late life social activity (76), purpose in life (2), well-being (74, 75) and time horizon. Time horizon was measured using 6 items (“1. Many opportunities await me in the future.”, “2. I expect that I will set new goals in the future.”, “ 3. I have the sense that time is running out.”, “4. I can do anything I want in the future.”, “5. There are limited possibilities in my future.”, “6. As I get older, I experience time as more limited.”) for which participants were asked to rate how each item applies to themselves using a 7-point Likert rating scale (Strongly agree (1), Agree (2), slightly agree (3), Neither agree nor disagree (4), Slightly disagree (5), Disagree (6), Strongly disagree (7)). Items (1,2 and 4) that are positively worded are flipped so that higher ratings on all items indicate a longer perceived time horizon. The total score is the mean of the item ratings, with a higher score indicating a longer perceived time horizon. Negative psychosocial exposures measured annually included negative life events (111), perceived social isolation (8), depressive symptoms (112), negative mood (113) and perceived stress (114). Adverse childhood experiences (115) were measured once. Procedure to request data and detailed information on the questionnaires can be found on the Rush Alzheimer’s Disease Center (RADC) Research Resource Sharing Hub website: https://www.radc.rush.edu/.

#### ROSMAP psychosocial variables summary scores

Each questionnaire score was transformed into a z-score (x-mean/sd) before being transformed into a t-score with an average of 100 and a standard deviation of 10 ((x*10)+100). Before computing the summary scores, the longitudinal variables were averaged across the follow-up. An index of positive exposures and an index of negative exposures were computed by averaging the t-scores for the positive and negative questionnaire scores respectively. The positive and negative scores included at least one assessment for each questionnaire. To create a global score that includes overall psychosocial experiences, positive variables were reversed and used to compute a combined negative score (social network size variable was excluded).

#### BLSA

For the BLSA cohort, summary scores of positive and negative psychosocial exposures were derived from the Activities and Attitudes Questionnaire (116). Detailed descriptions of the items included and calculations are provided in Supplemental doc 1. Depressive symptoms were assessed using the Center for Epidemiologic Studies Depression Scale (CESD)(117). Additionally, depressive and anxiety symptoms were also assessed using the Cornell Medical Index (118). For all the variables, the average of the values across the follow-up were used.

### Brain mitochondrial indices

#### ROSMAP proteomics

##### *A.* Tandem mass tag (TMT) isobaric labeling mass spectrometry

Post-mortem DLPFC (Broadman area 9) proteomics data measured with tandem mass tag (TMT) isobaric labeling mass spectrometry (66) was available for 400 (70% women) individuals including 168 cognitively normal individuals, 101 MCI, 123 AD and 8 had other causes of dementia. A total of 12,691 unique proteins were detected; 7,901 were kept after quality control.

Briefly, protein digestion was performed after tissue homogenization as previously described in (119). An equal amount of protein from each sample was aliquoted and digested in parallel to be used as global pooled internal standard (GIS) in the TMT batches. Before TMT labeling, sample randomization was performed by co-variates (age, sex, PMI, diagnosis, etc.) into 50 batches (8 cases per batch). The TMT 10-plex kit (ThermoFisher 90406) was used to label the samples (N=400) and the pooled GIS standards (N=100) as previously described (119, 120). In each batch, the 8 middle TMT channels were used to label individual samples and the TMT channels 126 and 131 were used to label GIS standards. High-pH fractionation was performed on an Agilent 1100 HPLC system as previously described (66, 121)). Peptide eluents separation was done on a self-packed C18 (1.9 μm, Dr. Maisch, Germany) fused silica column (25 cm × 75 μM internal diameter (ID); New Objective, Woburn, MA) by a Dionex UltiMate 3000 RSLCnano liquid chromatography system (ThermoFisher Scientific) and monitored using an Orbitrap Fusion mass spectrometer (ThermoFisher Scientific). The Proteome Discoverer software (version 2.3, ThermoFisher Scientific) was used to analyze the RAW files. The protein abundances were log2 transformed and regressed for batch, study, age at death and postmortem interval, prior to downstream analyses. The data is available through the synapse.org AMP-AD Knowledge Portal (www.synapse.org; SynapseID: syn17015098).

##### *B.* Selection reaction monitoring (SRM) quantitative proteomics

A small set of targeted proteins (N=119) were measured using selection reaction monitoring (SRM) quantitative proteomics as described previously (122) in post-mortem DLPFC brain tissue of 381 cognitively normal individuals, 286 MCI, 519 AD and 22 with other cause of dementia (68% women). The data is available through the synapse.org AMP-AD Knowledge Portal (www.synapse.org; SynapseID: syn10468856).

#### ROSMAP RNA-seq

RNA sequencing was performed from DLPFC (N = 1102), PCC (N = 661) AC (N = 731) tissues as previously described (123, 124). Briefly, samples were extracted using Qiagen’s miRNeasy mini kit (cat. no. 217004) and the RNase free DNase Set (cat. no. 79254). Quantification by Nanodrop and quality evaluation was performed by Agilent Bioanalyzer. The Illumina HiSeq with 101-bp paired-end reads was used for sequencing. To pass quality control, the mean coverage for samples was set at 95 million reads (median 90 million reads). Data included in the analysis met quality control criteria. A normalized expression level was computed for each gene by subtracting the mean expression across all samples and dividing by the SD. Before downstream analysis, the expression levels were regressed for batch, library size, percentage of coding bases, percentage of aligned reads, percentage of ribosomal bases, percentage of UTR base, median 5 prime to 3 prime base, median CV coverage, study (ROS or MAP) and post-mortem interval (PMI). The data is available through the synapse.org AMP-AD Knowledge Portal (www.synapse.org; SynapseID: syn3388564).

#### ROSMAP single-nucleus pseudo-bulk RNAseq data

Single-nucleus RNAseq was performed from frozen DLPFC specimens (N=424) as described previously (78, 125). Gray matter was extracted and dissociated into nuclei suspension. The Chromium Single Cell 3’ Reagent Kits version 3 (10x Genomics) were used to construct the single-nucleus RNA-seq libraries following the manufacturer’s protocol. The sample single-nucleus RNA-seq libraries were then sequenced using HiSeqX and NovaSeq sequencers (Illumina). CellRanger (v6.0.0; 10x Genomics) with the “GRCh38-2020-A” transcriptome and the “include-introns” option were used to process FASTQ files. The CellBender software was used for Cell calling and ambient RNA removal.

Cell clustering was performed based on existing cell type annotations (126) as described previously (125). A stepwise clustering approach was used to first identify 7 major cell types which were then subdivided into 92 cell subtypes. Pseudo-bulk UMI count matrices were constructed for each cell type and cell subtype by extracting and aggregating UMI counts of the cell (sub)type of interest for each participant, and normalizing them by sequencing depth. Pseudo-bulk UMI counts normalization was done by using the trimmed mean of M-values (TMM) method of edgeR, and log2 of counts per million174 mapped reads (CPM) were calculated using the voom function of limma (version 3.44.3). Low expression genes (log2CPM<2.0) were filtered out. Expression levels were quantile normalized after batch effects correction using ComBat. We selected cell types detected for >50% of participants and which had coverage for >50% of mitochondrial genes to compute the *MitoPathways* scores. Values were regressed for batch, PMI, age at death, study, sex, and cognitive status prior analysis. The raw and pseudo-bulk data is available through the AD Knowledge Portal (https://www.synapse.org/#!Synapse:syn31512863).

#### ROSMAP WGS mtDNAcn

WGS libraries were prepared using the KAPA Hyper Library Preparation Kit in accordance with the manufacturer’s instructions. Briefly, 650ng of DNA from DLPFC tissue was sheared using a Covaris LE220 sonicator (adaptive focused acoustics). After bead-based size selection, selected DNA fragments were end-repaired, adenylated, and ligated to Illumina sequencing adapters. To evaluate final libraries, fluorescent-based assays were used including qPCR with the Universal KAPA Library Quantification Kit and Fragment Analyzer (Advanced Analytics) or BioAnalyzer (Agilent 2100). Then, libraries were sequenced on an Illumina HiSeq X sequencer (v2.5 chemistry) using 2 x 150bp cycles. R/Bioconductor (packages GenomicAlignments and GenomicRanges) was used to calculate the median sequence coverages of the autosomal chromosomes and of the mitochondrial genome. Ambiguous regions were excluded using the intra-contig ambiguity mask from the BSgenome package. The mtDNAcn was calculated as (covmt/covnuc) × 2. Before downstream analysis, mtDNAcn was z-standardized and logarithmized as described previously (127). The data is available through the synapse.org AMP-AD Knowledge Portal (www.synapse.org; SynapseID: syn25618990).

#### BLSA proteomics

TMT proteomics was measured on post-mortem samples from 47 participants from two brain regions (middle frontal gyrus and precuneus). Detailed description of the method is provided in Supplemental doc 2. Values were regressed for age at death, PMI, sex and AD status before analysis.

#### Mitochondrial pathways’ indices calculation

Each datapoint (protein or transcript level) was transformed into a z-score ((x-mean)/sd) before being transformed into a t-score with an average of 100 and a standard deviation of 10 ((x*10)+100. Then for each score, t-scores (protein or transcript levels) were averaged to build a score representing the average abundance for each mitochondrial pathway (*MitoPathway*s) category. The *MitoPathway*s were defined using MitoCarta 3.0. mitochondrial gene annotations (73) (for study coverage see Table. S2). An estimated mitochondrial content index was calculated using the average protein expression of all mitochondrial genes referenced in MitoCarta 3.0. OxPhos enrichment scores were computed by calculating the ratio of each OxPhos category divided by the mitochondrial content score. Mitochondrial DNA density was computed using the ratio of mtDNA copies per cell (mtDNAcn derived from WGS) relative to mitochondrial content per cell (mito content index) as described previously (128). To ensure our findings were not confounded by variation of cellular content, we investigated the relationship between a set of housekeeping proteins (derived from (129)) and psychosocial scores and found no evidence of an association (Fig. 1D) Finally, since both OxPhos and mitochondrial mass were associated with psychosocial scores (Fig. 1D), the ratio of OxPhos protein levels to mitochondrial content score was computed to obtain a score representing specific mitochondrial OxPhos enrichment. To further increase the specificity of our analyses, we computed separate scores for subunits encoded by nuclear DNA (nDNA) and mtDNA.

### Statistical analysis

Protein and transcript levels were regressed for technical variables (see above) and post-mortal interval (PMI) prior to analysis. The associations between psychosocial scores and mitochondrial biology indices were assessed using Spearman’s rho correlation. Multivariate linear regressions were used to measure the relationship between psychosocial scores and respiratory complex I nDNA-encoded protein abundance scores adjusted for mitochondrial content while controlling for the effect of sex and cognitive status. We used principal component analysis (PCA) to visualize major cell type clustering after excluding genes with low expression across the entire dataset. Statistical analyses were performed with Prism 9 (GraphPad, CA) and RStudio version 1.4, and R version 4.0.4. Statistical significance was set at p<0.05.

## Supporting information

Supplemental doc 1

Supplemental doc 2

## Acknowledgments

We acknowledge participants of the ROS and MAP and BLSA studies. Support for this work was provided by NIH grants NHI RF1 AG057473 (PLD, DAB), U01 AG061356 (PLD, DAB), P30AG16101 (DAB), R01AG15819 (DAB), R01 AG070438 (PLD), R01AG17917 (DAB), U01AG46152 (PLD, DAB), U01AG61356 (PLD, DAB), R01AG036836 (PLD), U01AG061357 (SNT), R01AG061800 (SNT), RF1AG062181 (SNT), RF1AG076821 (MP), GM119793 (MP) and MH122706 (MP), the Intramural Program of the National Institute on Aging (MT), a NARSAD young investigator award (CT), the Nathaniel Wharton Fund and The Baszucki Brain (MP).

## Author contributions

Caroline Trumpff and Martin Picard designed the study. Vilas Menon, Hans-Ulrich Klein, Annie Lee, Vladislav Petyuk, Masashi Fujita, Cheyenne Hurst, Duc M. Duong, Nicholas T. Seyfried, Aliza Wingo, Thomas Wingo, Madhav Thambisetty contributed analysis tools or data. Caroline Trumpff and Martin Picard designed the analytic plan, and Caroline Trumpff and Anna Monzel performed the data analysis. Caroline Trumpff and Martin Picard drafted the original draft of the manuscript. All authors participated in the writing, reviewing and editing of the manuscript.

## References

1. B. S. McEwen, Physiology and neurobiology of stress and adaptation: central role of the brain. Physiological reviews 87, 873–904 (2007).

2. P. A. Boyle, L. L. Barnes, A. S. Buchman, D. A. Bennett, Purpose in life is associated with mortality among community-dwelling older persons. Psychosomatic medicine 71, 574 (2009).

3. P. A. Boyle, A. S. Buchman, L. L. Barnes, D. A. Bennett, Effect of a purpose in life on risk of incident Alzheimer disease and mild cognitive impairment in community-dwelling older persons. Archives of general psychiatry 67, 304–310 (2010).

4. T. E. Seeman, T. M. Lusignolo, M. Albert, L. Berkman, Social relationships, social support, and patterns of cognitive aging in healthy, high-functioning older adults: MacArthur studies of successful aging. Health psychology 20, 243 (2001).

5. L. Fratiglioni, H.-X. Wang, K. Ericsson, M. Maytan, B. Winblad, Influence of social network on occurrence of dementia: a community-based longitudinal study. The lancet 355, 1315–1319 (2000).

6. T. E. Seeman, G. A. Kaplan, L. Knudsen, R. Cohen, J. Guralnik, Social network ties and mortality among tile elderly in the Alameda County Study. American journal of epidemiology 126, 714–723 (1987).

7. J. T. Cacioppo, L. C. Hawkley, Perceived social isolation and cognition. *Trends in cognitive sciences* **13**, 447–454 (2009).

8. R. S. Wilson et al., Loneliness and risk of Alzheimer disease. Archives of general psychiatry 64, 234–240 (2007).

9. A. Terracciano et al., Personality and risk of Alzheimer’s disease: new data and meta-analysis. Alzheimer’s & Dementia 10, 179–186 (2014).

10. M. Picard, B. S. McEwen, Psychological stress and mitochondria: A systematic review. Psychosom Med 80, 141–153 (2018).

11. M. Picard, B. S. McEwen, Psychological stress and mitochondria: a conceptual framework. Psychosomatic medicine 80, 126 (2018).

12. M. Picard, C. Trumpff, Y. Burelle, Mitochondrial psychobiology: foundations and applications. *Current Opinion in Behavioral Sciences* **28**, 142–151 (2019).

13. T. E. Daniels, E. M. Olsen, A. R. Tyrka, Stress and Psychiatric Disorders: The Role of Mitochondria. Annual Review of Clinical Psychology 16 (2020).

14. J. Allen, H. J. Caruncho, L. E. Kalynchuk, Severe life stress, mitochondrial dysfunction, and depressive behavior: A pathophysiological and therapeutic perspective. Mitochondrion 56, 111–117 (2021).

15. Y. Chen, J. Zhang, How Energy Supports Our Brain to Yield Consciousness: Insights From Neuroimaging Based on the Neuroenergetics Hypothesis. Frontiers in Systems Neuroscience, 56 (2021).

16. P. J. Magistretti, I. Allaman, A cellular perspective on brain energy metabolism and functional imaging. Neuron 86, 883–901 (2015).

17. D. E. Befroy, K. F. Petersen, D. L. Rothman, G. I. Shulman, Assessment of in vivo mitochondrial metabolism by magnetic resonance spectroscopy. Methods in enzymology 457, 373–393 (2009).

18. Y. Ren et al., The impact of loneliness and social isolation on the development of cognitive decline and Alzheimer’s Disease. Frontiers in Neuroendocrinology, 101061 (2023).

19. Y. Yu et al., A 3D atlas of functional human brain energetic connectome based on neuropil distribution. Cerebral Cortex 33, 3996–4012 (2023).

20. S. J. Lupien et al., Cortisol levels during human aging predict hippocampal atrophy and memory deficits. Nature neuroscience 1, 69–73 (1998).

21. A. N. Clausen et al., Combat exposure, posttraumatic stress disorder, and head injuries differentially relate to alterations in cortical thickness in military veterans. Neuropsychopharmacology 45, 491–498 (2020).

22. V. Rangaraju et al., Pleiotropic mitochondria: the influence of mitochondria on neuronal development and disease. Journal of Neuroscience 39, 8200–8208 (2019).

23. C. Batandier et al., Acute stress delays brain mitochondrial permeability transition pore opening. Journal of neurochemistry 131, 314–322 (2014).

24. Y. Kambe, A. Miyata, Potential involvement of the mitochondrial unfolded protein response in depressive-like symptoms in mice. Neuroscience letters 588, 166–171 (2015).

25. W. Liu, C. Zhou, Corticosterone reduces brain mitochondrial function and expression of mitofusin, BDNF in depression-like rodents regardless of exercise preconditioning. Psychoneuroendocrinology 37, 1057–1070 (2012).

26. R. G. Hunter et al., Stress and corticosteroids regulate rat hippocampal mitochondrial DNA gene expression via the glucocorticoid receptor. Proc Natl Acad Sci U S A 113, 9099–9104 (2016).

27. M. Weger et al., Mitochondrial gene signature in the prefrontal cortex for differential susceptibility to chronic stress. Scientific reports 10, 18308 (2020).

28. S. Chakravarty et al., Chronic unpredictable stress (CUS)-induced anxiety and related mood disorders in a zebrafish model: altered brain proteome profile implicates mitochondrial dysfunction. PloS one 8, e63302 (2013).

29. M. D. Filiou et al., Proteomics and metabolomics analysis of a trait anxiety mouse model reveals divergent mitochondrial pathways. Biological psychiatry 70, 1074–1082 (2011).

30. A. R. Tyrka et al., Alterations of Mitochondrial DNA Copy Number and Telomere Length with Early Adversity and Psychopathology. Biol Psychiatry 10.1016/j.biopsych.2014.12.025 (2015).

31. N. Cai et al., Molecular signatures of major depression. Curr Biol 25, 1146–1156 (2015).

32. M. Picard et al., A mitochondrial health index sensitive to mood and caregiving stress. Biological psychiatry (2018).

33. M. Zvěřová et al., Disturbances of mitochondrial parameters to distinguish patients with depressive episode of bipolar disorder and major depressive disorder. Neuropsychiatric disease and treatment, 233–240 (2019).

34. C. Boeck et al., The association between cortisol, oxytocin and immune cell mitochondrial oxygen consumption in postpartum women with childhood maltreatment. Psychoneuroendocrinology (2018).

35. A. Karabatsiakis et al., Mitochondrial respiration in peripheral blood mononuclear cells correlates with depressive subsymptoms and severity of major depression. Transl Psychiatry 4, e397 (2014).

36. C. Boeck et al., Targeting the association between telomere length and immuno-cellular bioenergetics in female patients with Major Depressive Disorder. Scientific reports 8, 9419 (2018).

37. A. M. Gumpp et al., Childhood maltreatment is associated with changes in mitochondrial bioenergetics in maternal, but not in neonatal immune cells. Proceedings of the National Academy of Sciences 117, 24778–24784 (2020).

38. J. Hroudová, Z. Fišar, E. Kitzlerová, M. Zvěřová, J. Raboch, Mitochondrial respiration in blood platelets of depressive patients. Mitochondrion 13, 795–800 (2013).

39. M. Picard et al., Mitochondrial functions modulate neuroendocrine, metabolic, inflammatory, and transcriptional responses to acute psychological stress. Proceedings of the National Academy of Sciences 112, E6614–E6623 (2015).

40. U. Gimsa, E. Kanitz, W. Otten, S. M. Ibrahim, Behavior and stress reactivity in mouse strains with mitochondrial DNA variations. Annals of the New York Academy of Sciences 1153, 131–138 (2009).

41. T. L. Emmerzaal et al., Impaired mitochondrial complex I function as a candidate driver in the biological stress response and a concomitant stress-induced brain metabolic reprogramming in male mice. Translational psychiatry 10, 176 (2020).

42. T. Kasahara et al., Mice with neuron-specific accumulation of mitochondrial DNA mutations show mood disorder-like phenotypes. Molecular psychiatry 11, 577–593 (2006).

43. F. Hollis et al., Mitochondrial function in the brain links anxiety with social subordination. Proceedings of the National Academy of Sciences 112, 15486–15491 (2015).

44. M. A. van der Kooij et al., Diazepam actions in the VTA enhance social dominance and mitochondrial function in the nucleus accumbens by activation of dopamine D1 receptors. Molecular psychiatry 23, 569–578 (2018).

45. E. Gebara et al., Mitofusin-2 in the nucleus accumbens regulates anxiety and depression-like behaviors through mitochondrial and neuronal actions. Biological Psychiatry 89, 1033–1044 (2021).

46. X. Xie et al., Depression-like behaviors are accompanied by disrupted mitochondrial energy metabolism in chronic corticosterone-induced mice. The Journal of steroid biochemistry and molecular biology 200, 105607 (2020).

47. F. Duclot, L. Sailer, P. Koutakis, Z. Wang, M. Kabbaj, Transcriptomic regulations underlying pair-bond formation and maintenance in the socially monogamous male and female prairie vole. Biological Psychiatry 91, 141–151 (2022).

48. G. E. Ghosal S, Ramos-Fernández E, Chioino A, Grosse J, Guillot de Suduiraut I, Zanoletti O, Schneider B, Zorzano A, Astori S, Sandi C, Mitofusin-2 in nucleus accumbens D2-MSNs regulates social dominance and neuronal function. Cell reports 42(7):112776 (2023).

49. D. H. Ülgen, S. R. Ruigrok, C. Sandi, Powering the social brain: Mitochondria in social behaviour. Current Opinion in Neurobiology 79, 102675 (2023).

50. A. M. Rosenberg et al., A Network Approach to Mapping Mouse Brain-wide Mitochondrial Respiratory Chain Capacity in Relation to Behavior. Nature communications, 2021.2006. 2002.446767 (2023).

51. Y. Kato et al., Lower availability of mitochondrial complex I in anterior cingulate cortex in autism: a positron emission tomography study. American Journal of Psychiatry 180, 277–284 (2023).

52. L. Holper, D. Ben-Shachar, J. Mann, Multivariate meta-analyses of mitochondrial complex I and IV in major depressive disorder, bipolar disorder, schizophrenia, Alzheimer disease, and Parkinson disease. Neuropsychopharmacology 44, 837–849 (2019).

53. S. Rausser et al., Mitochondrial phenotypes in purified human immune cell subtypes and cell mixtures. Elife 10 (2021).

54. M. J. Devine, J. T. Kittler, Mitochondria at the neuronal presynapse in health and disease. Nature Reviews Neuroscience 19, 63–80 (2018).

55. R. Iwata, P. Vanderhaeghen, Regulatory roles of mitochondria and metabolism in neurogenesis. Current opinion in neurobiology 69, 231–240 (2021).

56. C. Fiebig et al., Mitochondrial dysfunction in astrocytes impairs the generation of reactive astrocytes and enhances neuronal cell death in the cortex upon photothrombotic lesion. Frontiers in molecular neuroscience 12, 40 (2019).

57. J. Ye et al., Electron transport chain inhibitors induce microglia activation through enhancing mitochondrial reactive oxygen species production. Experimental cell research 340, 315–326 (2016).

58. S. B. Shaikh, L. F. Nicholson, Effects of chronic low dose rotenone treatment on human microglial cells. Molecular Neurodegeneration 4, 1–13 (2009).

59. R. Schoenfeld et al., Oligodendroglial differentiation induces mitochondrial genes and inhibition of mitochondrial function represses oligodendroglial differentiation. Mitochondrion 10, 143–150 (2010).

60. A. Quintana, S. E. Kruse, R. P. Kapur, E. Sanz, R. D. Palmiter, Complex I deficiency due to loss of Ndufs4 in the brain results in progressive encephalopathy resembling Leigh syndrome. Proceedings of the National Academy of Sciences 107, 10996–11001 (2010).

61. A. Quintana et al., Fatal breathing dysfunction in a mouse model of Leigh syndrome. The Journal of clinical investigation 122, 2359–2368 (2012).

62. R. Iwata, P. Casimir, P. Vanderhaeghen, Mitochondrial dynamics in postmitotic cells regulate neurogenesis. Science 369, 858–862 (2020).

63. G. Sturm et al., OxPhos defects cause hypermetabolism and reduce lifespan in cells and in patients with mitochondrial diseases. Communications biology 6, 1–22 (2023).

64. R. H. Swerdlow, J. M. Burns, S. M. Khan, The Alzheimer’s disease mitochondrial cascade hypothesis: progress and perspectives. Biochimica et Biophysica Acta (BBA)-Molecular Basis of Disease 1842, 1219–1231 (2014).

65. D. A. Bennett et al., Religious orders study and rush memory and aging project. Journal of Alzheimer’s Disease 64, S161–S189 (2018).

66. C. Robins et al., Genetic control of the human brain proteome. bioRxiv, 816652 (2019).

67. K. N. Ochsner, J. J. Gross, The cognitive control of emotion. Trends in cognitive sciences 9, 242–249 (2005).

68. A. F. Arnsten, Stress weakens prefrontal networks: molecular insults to higher cognition. Nature neuroscience 18, 1376 (2015).

69. E. B. Ansell, K. Rando, K. Tuit, J. Guarnaccia, R. Sinha, Cumulative adversity and smaller gray matter volume in medial prefrontal, anterior cingulate, and insula regions. Biological psychiatry 72, 57–64 (2012).

70. C. Liston, B. S. McEwen, B. Casey, Psychosocial stress reversibly disrupts prefrontal processing and attentional control. Proceedings of the National Academy of Sciences 106, 912–917 (2009).

71. A. Monzel, J. Enriques, M. Picard, Multifaceted mitochondria: Moving mitochondrial science beyond function and dysfunction. *Nature Metabolism* In press (2023).

72. B. K. Chacko, D. Zhi, V. M. Darley-Usmar, T. Mitchell, The Bioenergetic Health Index is a sensitive measure of oxidative stress in human monocytes. Redox Biol 8, 43–50 (2016).

73. S. Rath et al., MitoCarta3. 0: an updated mitochondrial proteome now with sub-organelle localization and pathway annotations. *Nucleic Acids Research* (2020).

74. C. D. Ryff, C. L. M. Keyes, The structure of psychological well-being revisited. Journal of personality and social psychology 69, 719 (1995).

75. R. S. Wilson et al., The influence of cognitive decline on well-being in old age. Psychology and aging 28, 304 (2013).

76. A. S. Buchman et al., Association between late-life social activity and motor decline in older adults. Archives of internal medicine 169, 1139–1146 (2009).

77. C. Fecher et al., Cell-type-specific profiling of brain mitochondria reveals functional and molecular diversity. Nature neuroscience 22, 1731–1742 (2019).

78. G. S. Green et al., Cellular dynamics across aged human brains uncover a multicellular cascade leading to Alzheimer’s disease. bioRxiv, 2023.2003. 2007.531493 (2023).

79. I. Perić, V. Costina, A. Stanisavljević, P. Findeisen, D. Filipović, Proteomic characterization of hippocampus of chronically socially isolated rats treated with fluoxetine: Depression-like behaviour and fluoxetine mechanism of action. Neuropharmacology 135, 268–283 (2018).

80. G. T. Rezin et al., Inhibition of mitochondrial respiratory chain in brain of rats subjected to an experimental model of depression. Neurochemistry international 53, 395–400 (2008).

81. J. L. Madrigal et al., Glutathione depletion, lipid peroxidation and mitochondrial dysfunction are induced by chronic stress in rat brain. Neuropsychopharmacology 24, 420–429 (2001).

82. P. Rinwa, A. Kumar, Piperine potentiates the protective effects of curcumin against chronic unpredictable stress-induced cognitive impairment and oxidative damage in mice. Brain research 1488, 38–50 (2012).

83. P. Rinwa, A. Kumar, Modulation of nitrergic signalling pathway by American ginseng attenuates chronic unpredictable stress-induced cognitive impairment, neuroinflammation, and biochemical alterations. Naunyn-Schmiedeberg’s archives of pharmacology 387, 129–141 (2014).

84. E. Morava et al., Depressive behaviour in children diagnosed with a mitochondrial disorder. Mitochondrion 10, 528–533 (2010).

85. A. Strasser, L. Xin, R. Gruetter, C. Sandi, Nucleus accumbens neurochemistry in human anxiety: A 7 T 1H-MRS study. European neuropsychopharmacology 29, 365–375 (2019).

86. C. J. Jong, J. Azuma, S. Schaffer, Mechanism underlying the antioxidant activity of taurine: prevention of mitochondrial oxidant production. Amino acids 42, 2223–2232 (2012).

87. S. W. Schaffer, K. Shimada-Takaura, C. J. Jong, T. Ito, K. Takahashi, Impaired energy metabolism of the taurine-deficient heart. Amino Acids 48, 549–558 (2016).

88. A. Wiehler, F. Branzoli, I. Adanyeguh, F. Mochel, M. Pessiglione, A neuro-metabolic account of why daylong cognitive work alters the control of economic decisions. Current Biology 32, 3564–3575. e3565 (2022).

89. M. Picard et al., A mitochondrial health index sensitive to mood and caregiving stress. Biological psychiatry 84, 9–17 (2018).

90. C. Boeck et al., Inflammation in adult women with a history of child maltreatment: the involvement of mitochondrial alterations and oxidative stress. Mitochondrion 30, 197–207 (2016).

91. C. Boeck et al., The association between cortisol, oxytocin, and immune cell mitochondrial oxygen consumption in postpartum women with childhood maltreatment. Psychoneuroendocrinology 96, 69–77 (2018).

92. L. E. Gyllenhammer et al., Maternal inflammation during pregnancy and offspring brain development: the role of mitochondria. Biological Psychiatry: Cognitive Neuroscience and Neuroimaging 7, 498–509 (2022).

93. N. Bobba-Alves, R.-P. Juster, M. Picard, The energetic cost of allostasis and allostatic load. Psychoneuroendocrinology 146, 105951 (2022).

94. M. Schwaiger et al., Altered Stress-Induced Regulation of Genes in Monocytes in Adults with a History of Childhood Adversity. Neuropsychopharmacology 41, 2530–2540 (2016).

95. K. Chandramouli, P. Y. Qian, Proteomics: challenges, techniques and possibilities to overcome biological sample complexity. Hum Genomics Proteomics 2009 (2009).

96. L. E. Liharska et al., A study of gene expression in the living human brain. medRxiv, 2023.2004. 2021.23288916 (2023).

97. L. Yu et al., Cortical proteins associated with cognitive resilience in community-dwelling older persons. JAMA psychiatry 77, 1172–1180 (2020).

98. L. Ferrucci (2008) The Baltimore Longitudinal Study of Aging (BLSA): a 50-year-long journey and plans for the future. (Oxford University Press).

99. V. Swarup et al., Identification of conserved proteomic networks in neurodegenerative dementia. Cell Reports 31, 107807 (2020).

100. A. P. Wingo et al., Large-scale proteomic analysis of human brain identifies proteins associated with cognitive trajectory in advanced age. Nature communications 10, 1–14 (2019).

101. T. G. Beach et al., Arizona study of aging and neurodegenerative disorders and brain and body donation program. Neuropathology: official journal of the Japanese Society of Neuropathology 35, 354 (2015).

102. D. A Bennett, J. A Schneider, Z. Arvanitakis, R. S Wilson, Overview and findings from the religious orders study. Current Alzheimer Research 9, 628–645 (2012).

103. D. A Bennett et al., Overview and findings from the rush Memory and Aging Project. Current Alzheimer Research 9, 646–663 (2012).

104. D. Bennett et al., Neuropathology of older persons without cognitive impairment from two community-based studies. Neurology 66, 1837–1844 (2006).

105. D. A. Bennett et al., Natural history of mild cognitive impairment in older persons. Neurology 59, 198–205 (2002).

106. D. A. Bennett et al., Decision rules guiding the clinical diagnosis of Alzheimer’s disease in two community-based cohort studies compared to standard practice in a clinic-based cohort study. Neuroepidemiology 27, 169–176 (2006).

107. D. Bennett et al., Apolipoprotein E ε4 allele, AD pathology, and the clinical expression of Alzheimer’s disease. Neurology 60, 246–252 (2003).

108. B. T. Hyman, J. Q. Trojanowski, Editorial on consensus recommendations for the postmortem diagnosis of Alzheimer disease from the National Institute on Aging and the Reagan Institute Working Group on diagnostic criteria for the neuropathological assessment of Alzheimer disease. Journal of neuropathology and experimental neurology 56, 1095 (1997).

109. H.-U. Klein, M. Schäfer, D. A. Bennett, H. Schwender, P. L. De Jager, Bayesian integrative analysis of epigenomic and transcriptomic data identifies Alzheimer’s disease candidate genes and networks. PLoS computational biology 16, e1007771 (2020).

110. D. A. Bennett, J. A. Schneider, Y. Tang, S. E. Arnold, R. S. Wilson, The effect of social networks on the relation between Alzheimer’s disease pathology and level of cognitive function in old people: a longitudinal cohort study. The Lancet Neurology 5, 406–412 (2006).

111. R. S. Wilson et al., Negative social interactions and risk of mild cognitive impairment in old age. Neuropsychology 29, 561 (2015).

112. L. S. Radloff, The CES-D scale: A self-report depression scale for research in the general population. Applied psychological measurement 1, 385–401 (1977).

113. D. Watson, L. A. Clark, A. Tellegen, Development and validation of brief measures of positive and negative affect: the PANAS scales. Journal of personality and social psychology 54, 1063 (1988).

114. A. D. Turner, B. D. James, A. W. Capuano, N. T. Aggarwal, L. L. Barnes, Perceived stress and cognitive decline in different cognitive domains in a cohort of older African Americans. The American Journal of Geriatric Psychiatry 25, 25–34 (2017).

115. R. S. Wilson et al., Childhood adversity and psychosocial adjustment in old age. The American Journal of Geriatric Psychiatry 14, 307–315 (2006).

116. E. W. Burgess, R. S. Cavan, R. J. Havighurst, Your activities and attitudes. (1949).

117. W. W. Eaton, C. Smith, M. Ybarra, C. Muntaner, A. Tien, Center for Epidemiologic Studies Depression Scale: review and revision (CESD and CESD-R). (2004).

118. K. Brodman, A. J. Erdmann, I. Lorge, H. G. Wolff, T. H. BROADBENT, The Cornell Medical Index: an adjunct to medical interview. Journal of the American Medical Association 140, 530–534 (1949).

119. L. Ping et al., Global quantitative analysis of the human brain proteome in Alzheimer’s and Parkinson’s Disease. Scientific data 5, 1–12 (2018).

120. E. C. Johnson et al., Deep proteomic network analysis of Alzheimer’s disease brain reveals alterations in RNA binding proteins and RNA splicing associated with disease. Molecular neurodegeneration 13, 1–22 (2018).

121. P. Mertins et al., Reproducible workflow for multiplexed deep-scale proteome and phosphoproteome analysis of tumor tissues by liquid chromatography–mass spectrometry. Nature protocols 13, 1632–1661 (2018).

122. L. Yu et al., Targeted brain proteomics uncover multiple pathways to Alzheimer’s dementia. Annals of neurology 84, 78–88 (2018).

123. T. Raj et al., Integrative transcriptome analyses of the aging brain implicate altered splicing in Alzheimer’s disease susceptibility. Nature genetics 50, 1584–1592 (2018).

124. S. Oveisgharan et al., Estrogen receptor genes, cognitive decline, and Alzheimer disease. Neurology 100, e1474–e1487 (2023).

125. M. Fujita et al., Cell-subtype specific effects of genetic variation in the aging and Alzheimer cortex. bioRxiv, 2022.2011. 2007.515446 (2022).

126. A. Cain et al., Multicellular communities are perturbed in the aging human brain and Alzheimer’s disease. Nature Neuroscience, 1–14 (2023).

127. H.-U. Klein et al., Characterization of mitochondrial DNA quantity and quality in the human aged and Alzheimer’s disease brain. medRxiv 10.1101/2021.05.20.21257456, 2021.2005.2020.21257456 (2021).

128. C. Trumpff et al., Mitochondrial respiratory chain protein co-regulation in the human brain. Heliyon 8, e09353 (2022).

129. L. Jiang et al., A quantitative proteome map of the human body. Cell 183, 269–283. e219 (2020).

