## Supplemental figures for "Psychosocial experiences are associated with human brain mitochondrial biology"

### Supplementary figures

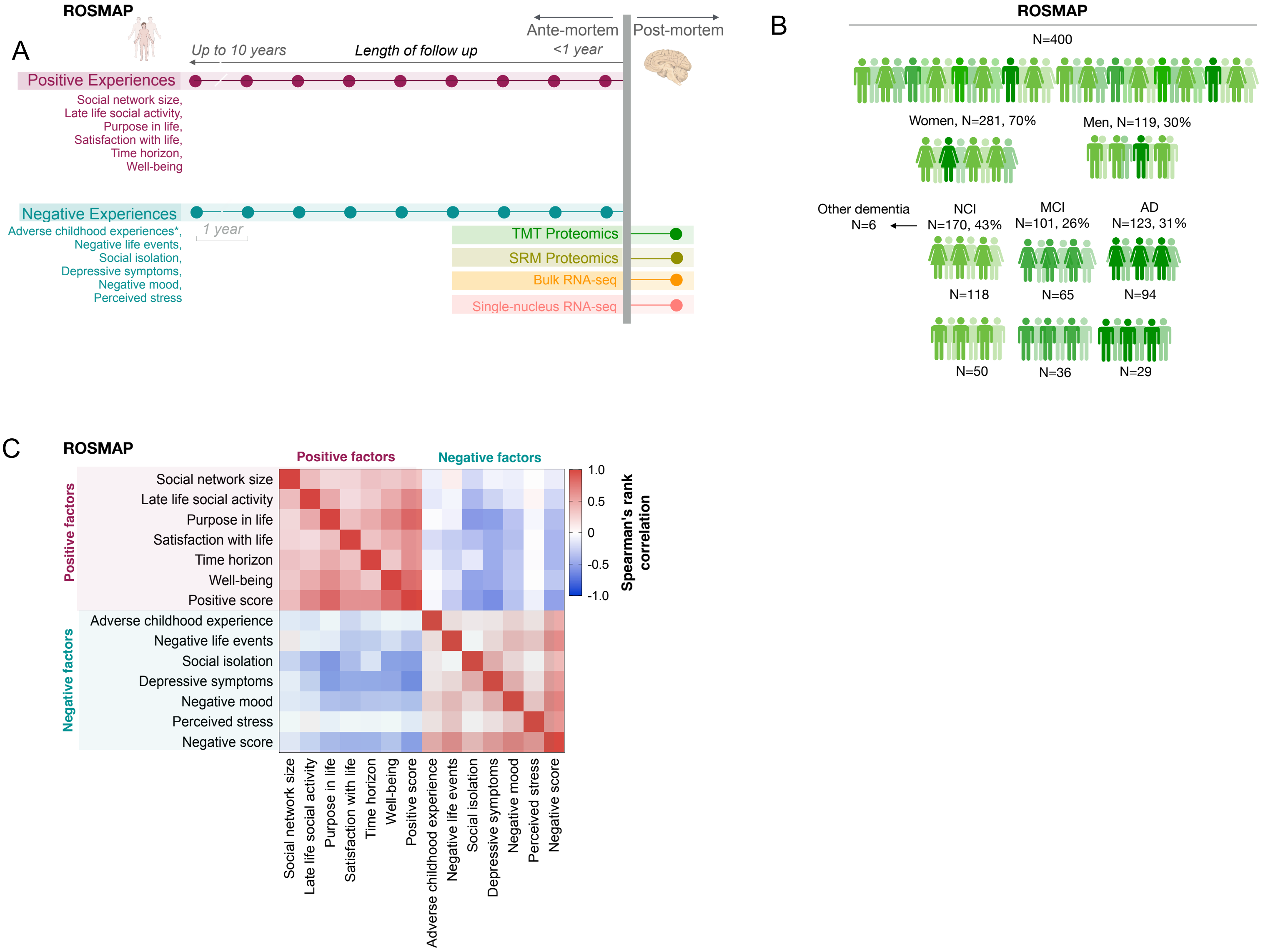


**Fig. S1. Religious Orders Study (ROS) and Rush Memory and Ageing Project (MAP) study design and psychosocial variables.** (**A**) Study design, (**B**) ROSMAP participant characteristics with post-mortem TMT proteomics data from the dorsolateral prefrontal cortex (N=400). Participant characteristics for other omics datasets are shown in Supplemental table S1. **(C)** Association between positive and negative psychosocial summary scores and individual questionnaire scores.

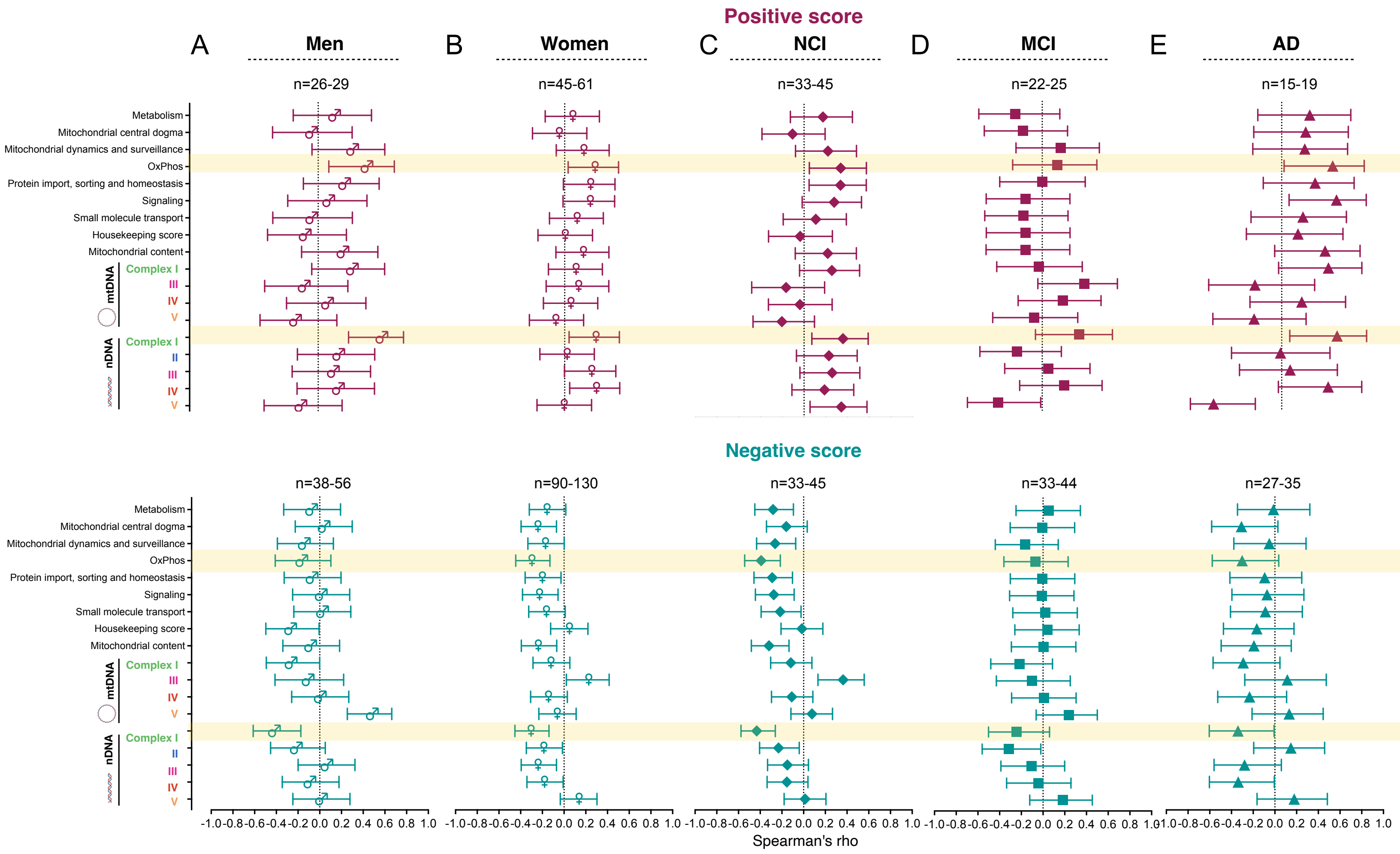


**Fig S2. Psychosocial factors and the mitochondrial brain proteome by sex and cognitive status (ROSMAP, TMT proteomics).** Since our sample included both men and women with various cognitive statuses by the time of death, we ran stratified analyses. Effect size (spearman's rho (95% CI)) for the association between positive and negative psychosocial scores and mitochondrial protein abundance (see Fig1) in (**A**) men, (**B**) women, individuals with (**C**) no cognitive impairment (NCI), (**D**) mild-cognitive impairment or (**E**) Alzheimer's disease (AD) at the time of death. Detailed results are shown in Supplemental table S3C. Protein abundances were regressed for age at death, postmortem interval, study and batch prior to analysis.

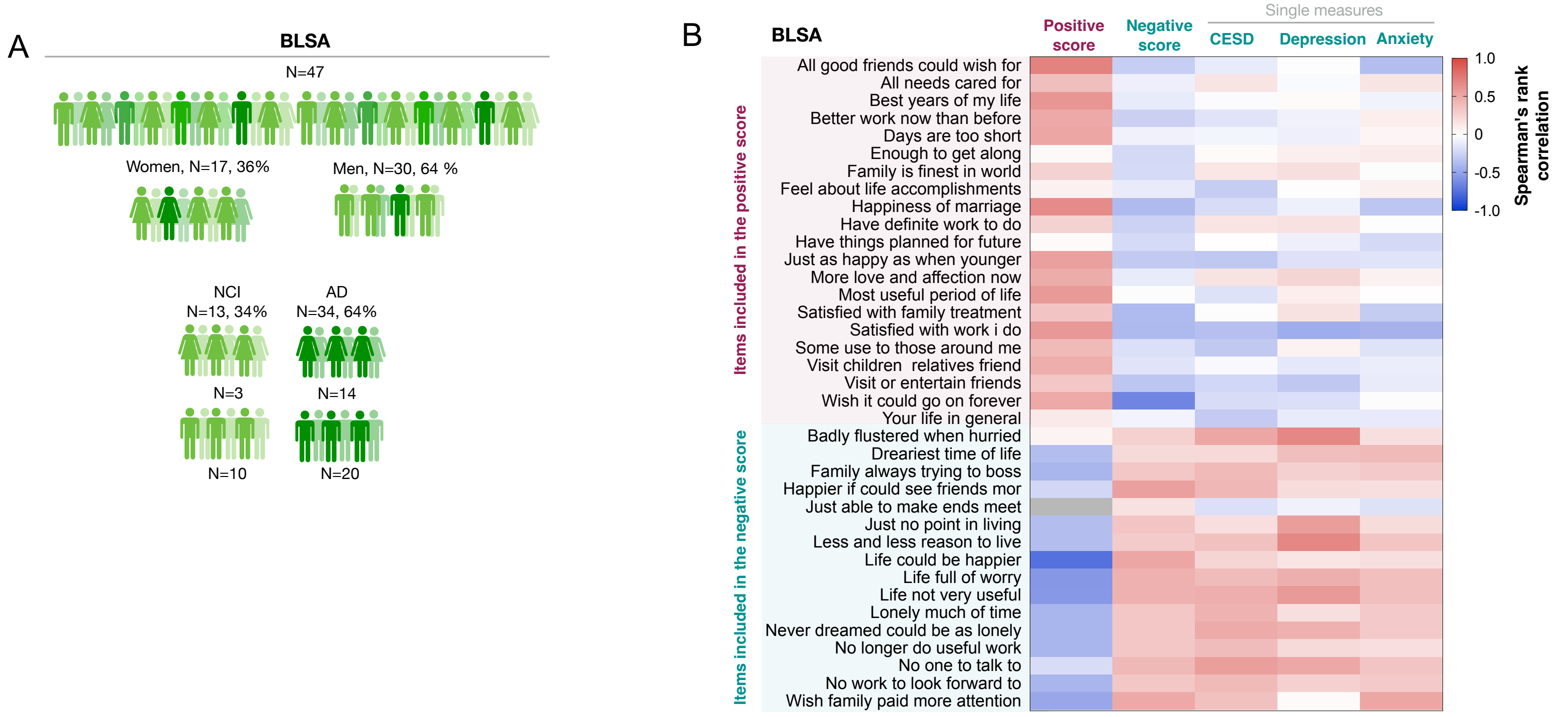


**Fig. S3. Baltimore Longitudinal Study of Aging (BLSA) participant characteristics and psychosocial variables and summary scores.** (**A**) BLSA participant characteristics with post-mortem data from the middle frontal gyrus. (**B**) Association between positive and negative psychosocial summary scores of the Activities and Attitude Questionnaires (Burgess et al., 1949). The individual item code was "agree = 1, disagree = 0" for all items excepted for items "happiness of marriage" = 1 = very unhappy, 2 = unhappy, 3 = average, 4 = happy, "Hear from young people" = 1 = once/year, 2= few times/year, 3=once or twice/month. 4=once/week, 5=every day, 0= have no young friends; "your life in general" = 1= very happy, 2 = moderately happy, 3 = average, 4 = unhappy; "feel about life accomplishments" = 1 = well satisfied, 2 = reasonably satisfied, 3 = dissatisfied. Depression and Anxiety scales were derived from the Cornell Medical Index questionnaire (Brodman et al., 1949)).

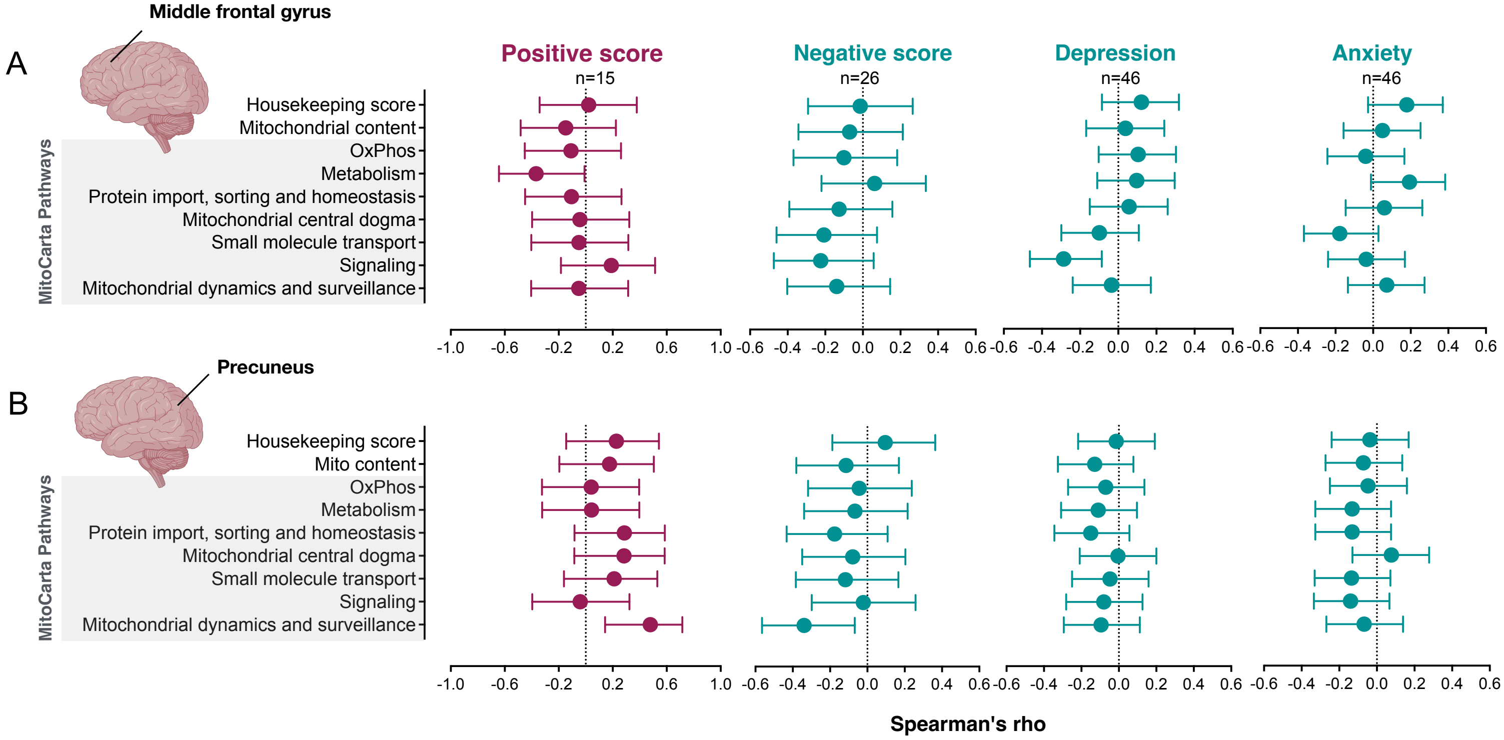


**Fig. S4. Psychosocial factors and mitochondrial protein abundance in BLSA.** Effect size (spearman's rho (95% CI)) for the association between psychosocial factors, housekeeping proteins and mitochondrial protein summary scores in (**A**) Middle frontal gyrus and the (**B**) Precuneus. For the study gene coverage see Supplemental table S2. Spearman's rho (95% CI). Scores were adjusted for post-mortem interval, age at death, sex and AD status. Detailed results are shown in table S3B.

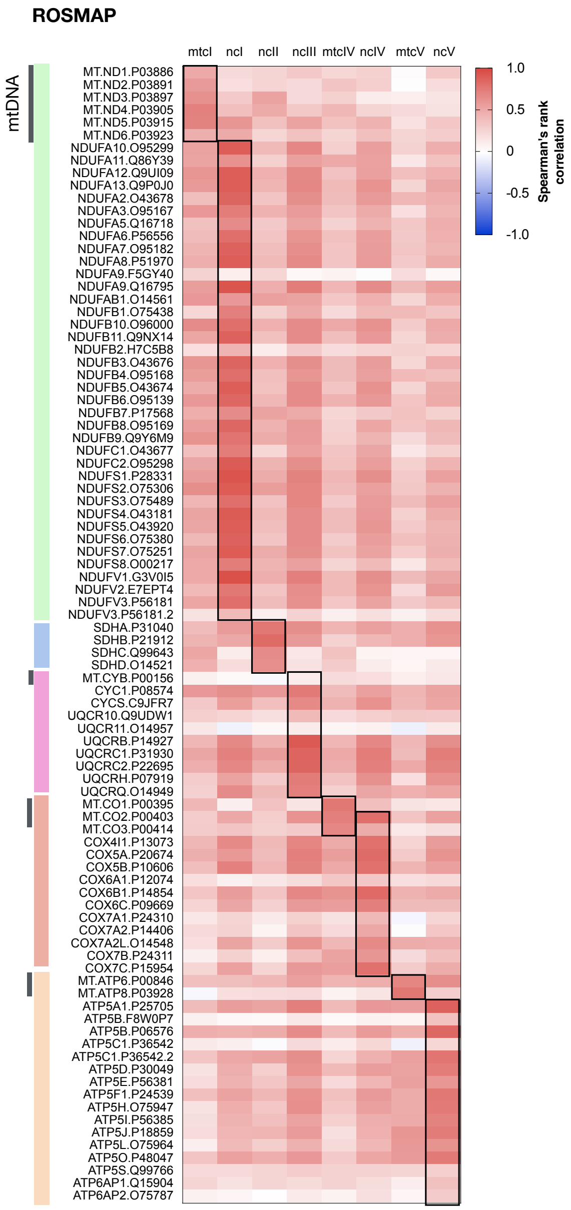


**Fig. S5. Correlation between mitochondrial OxPhos summary scores and individual proteins levels (ROSMAP).** Tandem mass tag (TMT) isobaric labeling mass spectrometry data on post-mortem dorsolateral prefrontal cortex (DLPFC). Results from Spearman rank correlations. N=400.

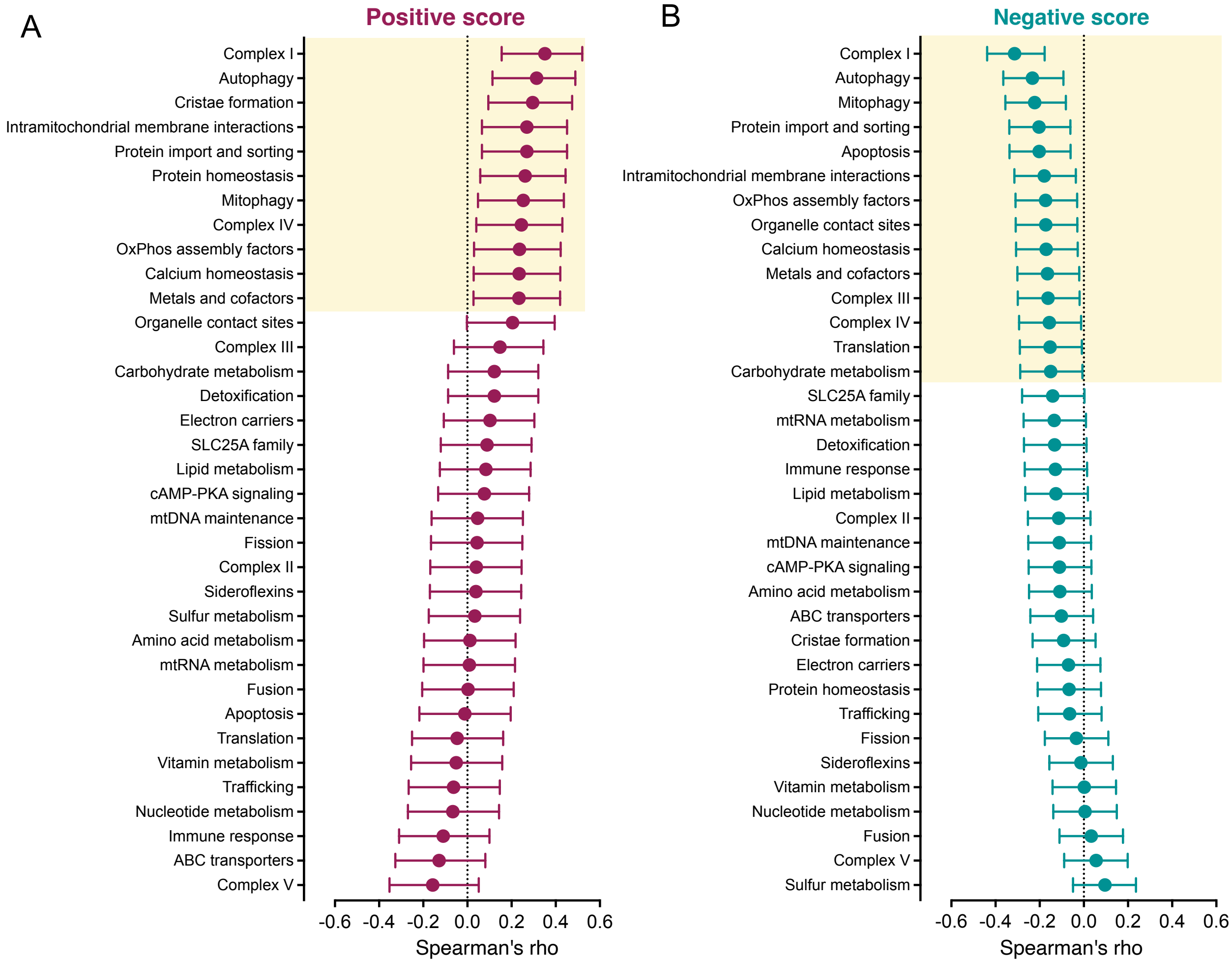


**Fig S6. Psychosocial factors and the mitochondrial brain proteome (ROSMAP, TMT, DLPFC): MitoCarta pathways levels 2 summary scores.** Effect size (spearman's rho (95% CI)) for the association between (A) positive and (B) negative psychosocial scores and MitoCarta 3.0. pathways levels 2. Protein abundances were regressed for age at death, postmortem interval, study and batch prior to analysis. N=400.

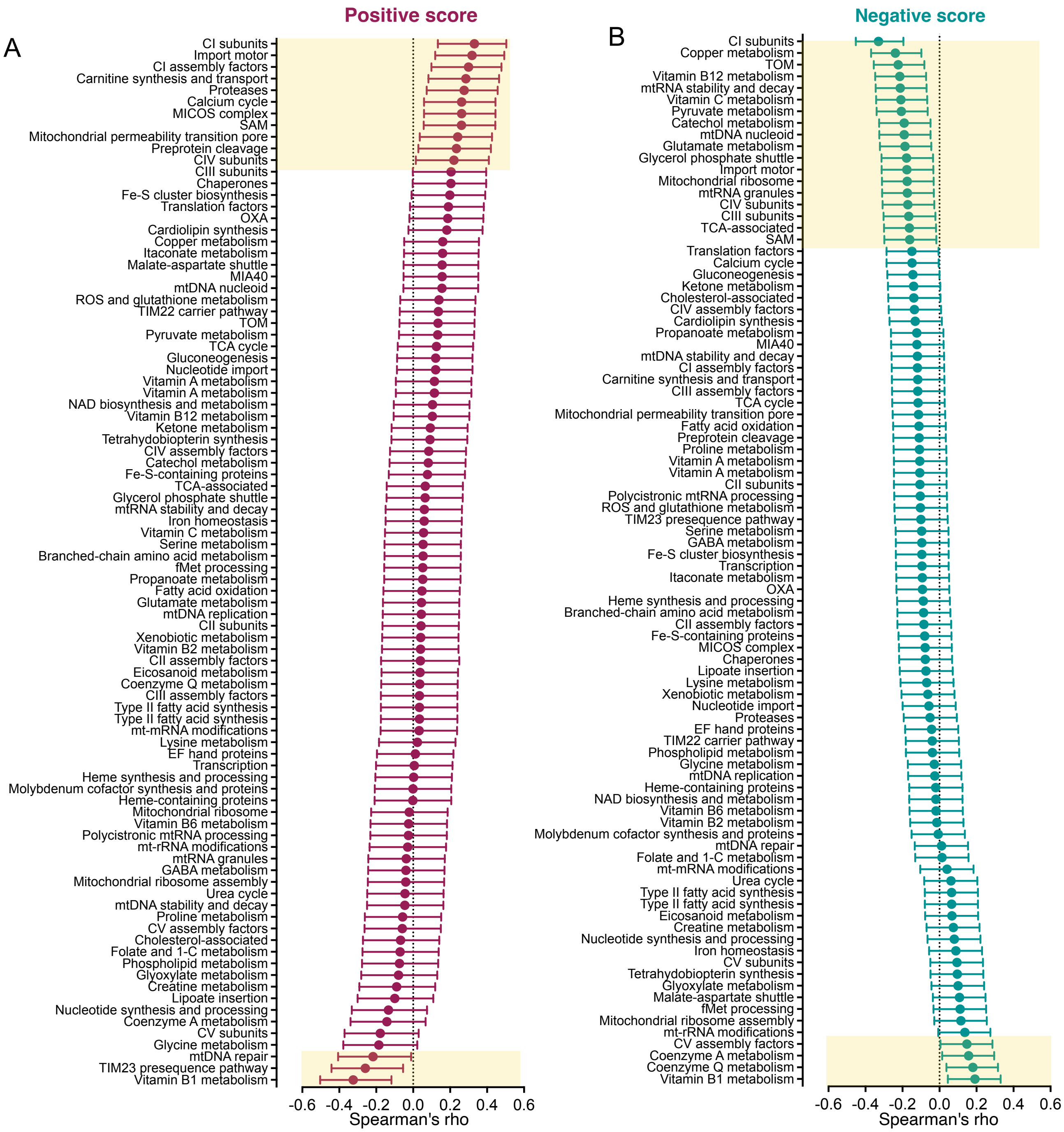


**Fig S7. Psychosocial factors and the mitochondrial brain proteome (ROSMAP): MitoCarta pathways levels 3 summary scores.** Effect size (spearman's rho (95% CI)) for the association between (A) positive and (B) negative psychosocial scores and MitoCarta 3.0. pathways levels 3. Protein abundances were regressed for age at death, postmortem interval, study and batch prior to analysis. N=400.

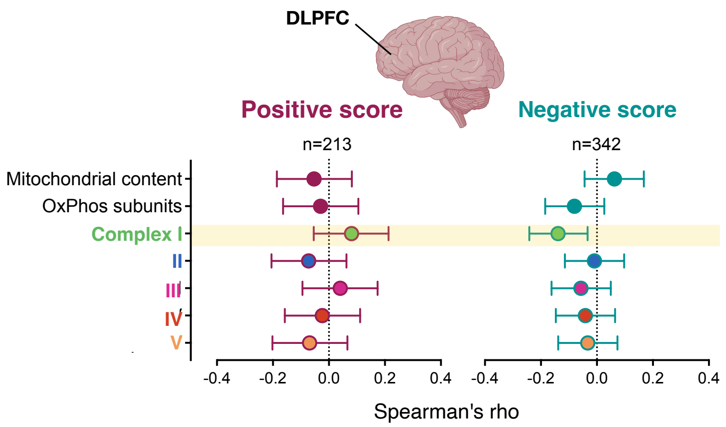


**Fig. S8. Psychosocial factors and the mitochondrial brain proteome (ROSMAP, SRM proteomics, N=1,208).** Effect size (spearman's rho (95% CI)) for the association between positive and negative psychosocial scores and mitochondrial protein abundance. Mitochondrial content score was derived from VDAC2. The OxPhos subunit score was composed by complex I (NDUFA10, NDUFA5, NDUFA6, NDUFA7, NDUFS6, NDUFV1), II (SDHA, SDHB, SDHC, SDHD), III (CYC1, UQCR10, UQCRC2), IV (COX7B) and V(ATP5F1, ATP5J2) proteins divided by the mitochondrial content score. Detailed results are shown in Supplemental table S3D.

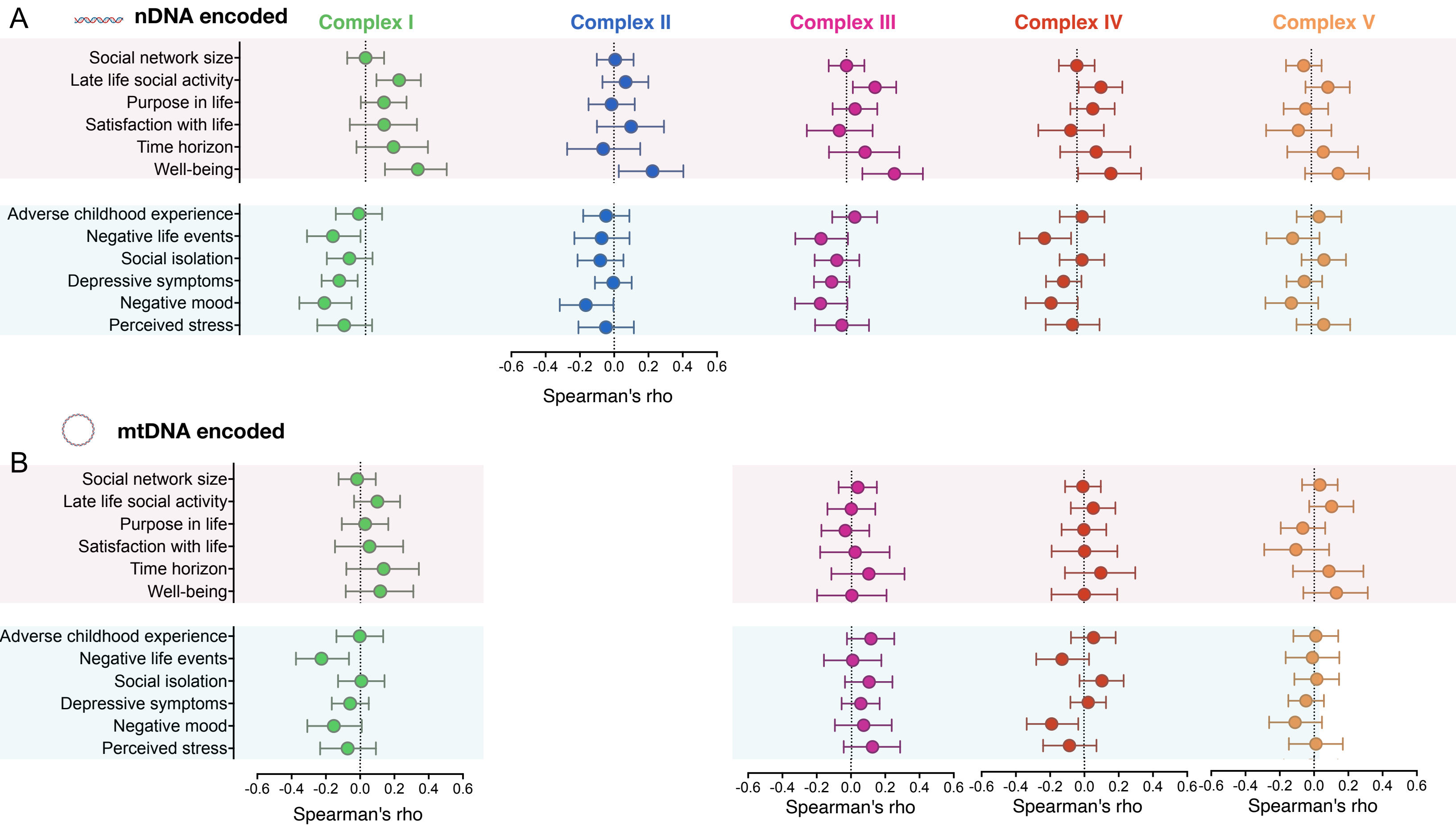


**Fig S9. Psychosocial factors and OxPhos brain protein abundance (ROSMAP, TMT, DLPFC): individual questionnaires.** Effect size (spearman's rho (95% CI)) for the association between psychosocial factors and (**A**) nuclear DNA and (**B**) mitochondrial DNA encoded mitochondrial OxPhos proteins. Detailed results are shown in Supplemental table S3F. Protein abundances were regressed for age at death, postmortem interval, study and batch prior to analysis.

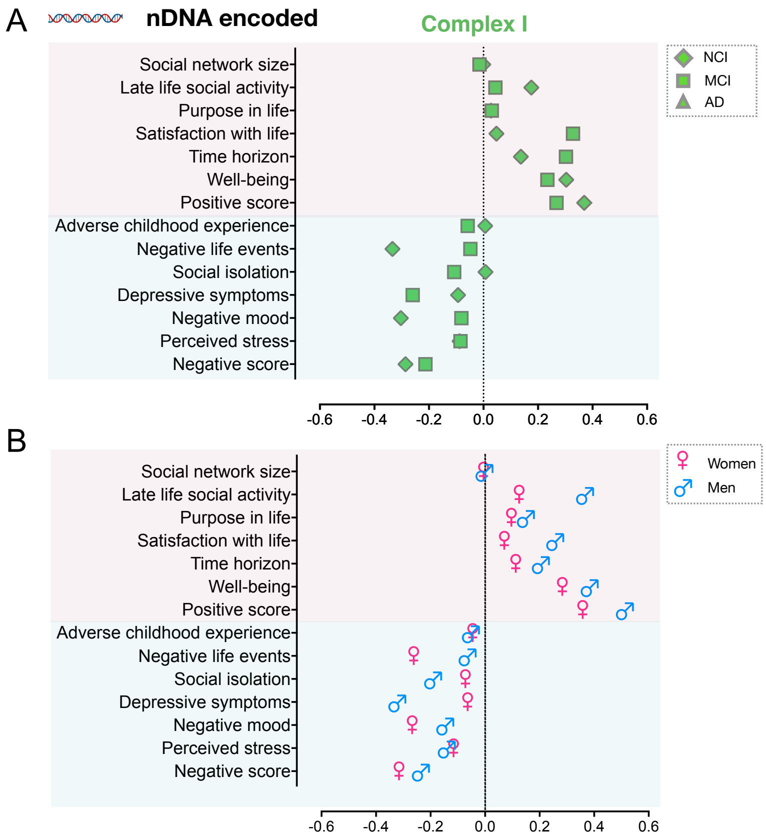


**Fig. S10. Psychosocial factors and mitochondrial respiratory chain complex I (nDNA-encoded) protein abundance (ROSMAP, TMT, DLPFC).** Analysis stratified by **(A)** cognitive status at the time of death and by **(B)** sex. Protein abundances were regressed for age at death, postmortem interval, study and batch prior to analysis. Positive score: NCI n=45, MCI n=25, AD n=19, men n=29, women n=61. Negative score: NCI n=104, MCI n=44, AD n=35, men n=56, women n=130.

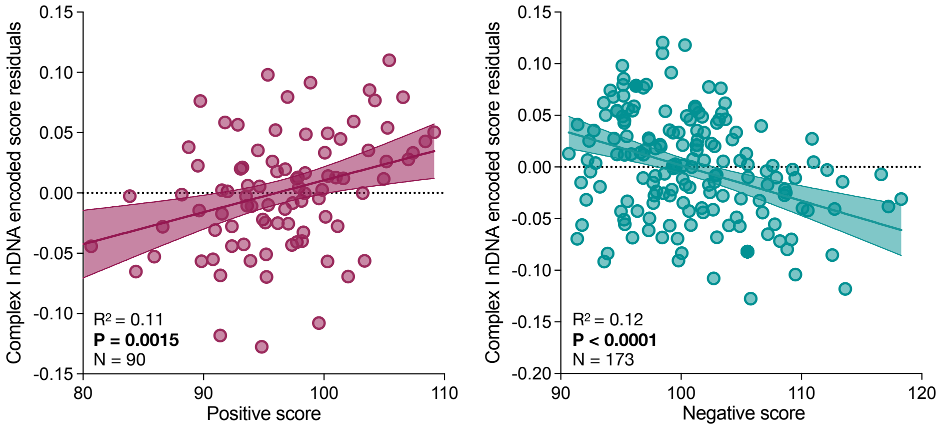


**Fig S11. Psychosocial factors and human brain mitochondrial respiratory chain complex I (nDNA-encoded) protein abundance.** Scatterplots of the relationship between (**A**) positive and (**B**) negative psychosocial scores and DLPFC complex I nDNA-encoded protein abundance adjusted for mitochondrial content. Results from multivariate linear regressions adjusting for sex and cognitive status. Protein abundances were regressed for age at death, postmortem interval, study and batch prior to analysis.

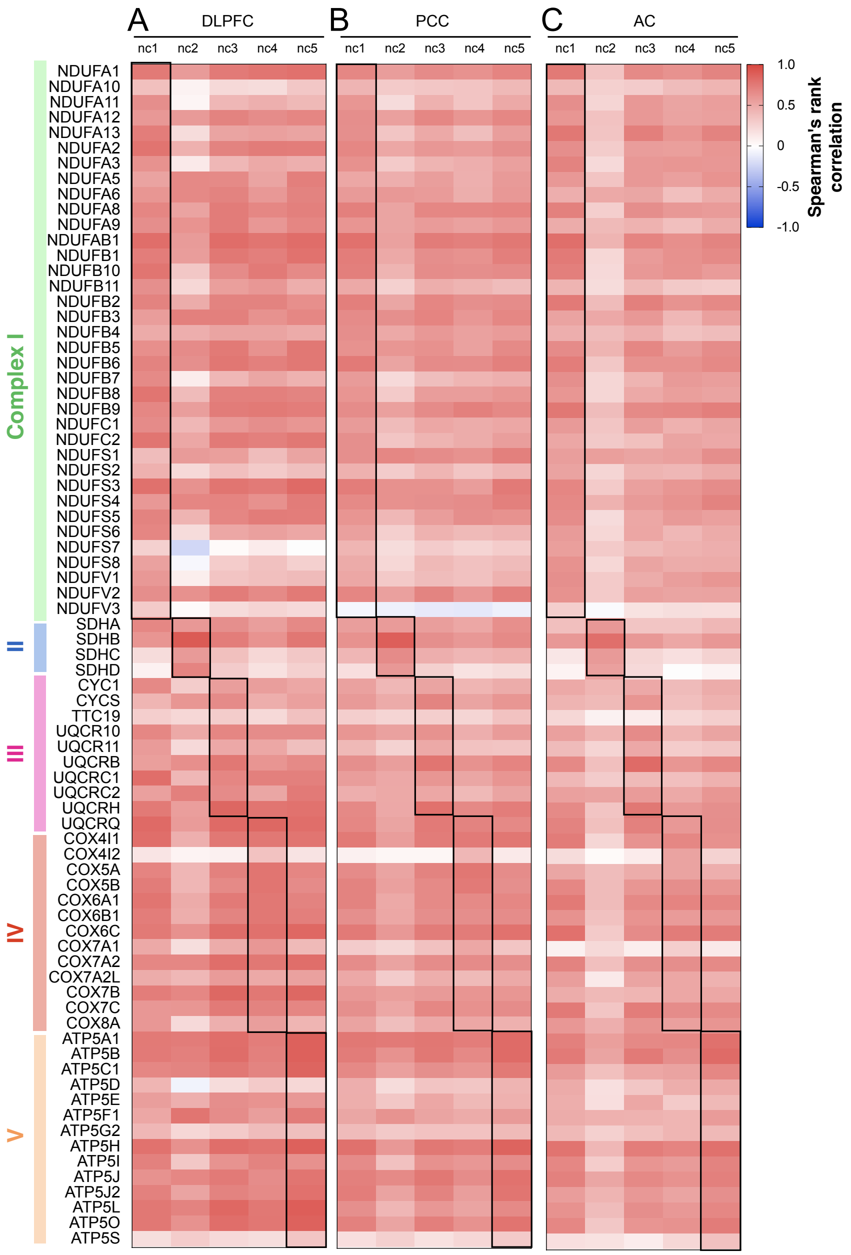


**Fig. S12. Correlation between mitochondrial OxPhos transcript summary scores and individual transcript levels (ROSMAP, RNA-sequencing).** Data from the (**A**) dorsolateral prefrontal cortex (DLPFC, N = 1102), the (**B**) posterior cingulate cortex (PCC, N = 661) and the (**C**) anterior caudate (AC, N = 731). Results from Spearman rank correlations.

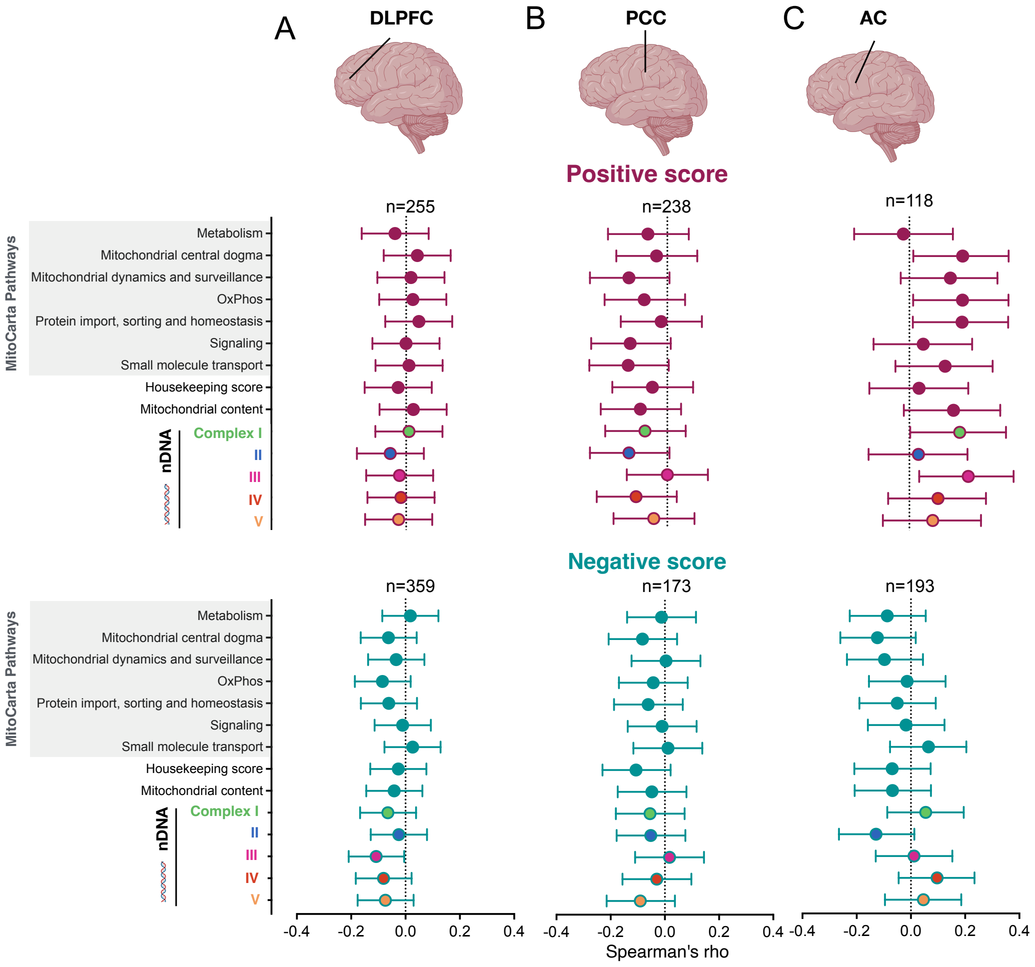


**Fig S13. Psychosocial factors and the mitochondrial brain transcriptome.** Effect size (spearman's rho (95% CI)) for the association between positive and negative psychosocial scores and mitochondrial transcript abundance (see Fig1) in the (**A**) dorsolateral prefrontal cortex (DLPFC), the (**B**) posterior cingulate cortex (PCC) and the (**C**) anterior caudate (AC). Detailed results are shown in Supplemental table S3E. Transcript abundances were regressed for batch, library size, percentage of coding bases, percentage of aligned reads, percentage of ribosomal bases, percentage of UTR base, median 5 prime to 3 prime base, median CV coverage, study (ROS or MAP) and post-mortem interval (PMI) prior to analysis.
