## Supplemental doc 1 for "Psychosocial experiences are associated with human brain mitochondrial biology"

**Activities and Attitudes Questionnaire Scoring**

List of items included in the final score:

|  | **Item label** | **Code** | **Score** |
| --- | --- | --- | --- |
| 1 | Have things planned for future | 0 = Not Checked, 1 = Checked | Positive sub-score 1 |
| 2 | Visit children, relatives, friends | 0 = Not Checked, 1 = Checked | Positive sub-score 1 |
| 3 | Visit or entertain friends | 0 = Not Checked, 1 = Checked | Positive sub-score 1 |
| 4 | All good friends could wish for | 1 = Agree, 0 = Disagree | Positive sub-score 2 |
| 5 | All needs cared for | 1 = Agree, 0 = Disagree | Positive sub-score 2 |
| 6 | Best years of my life | 1 = Agree, 0 = Disagree | Positive sub-score 2 |
| 7 | Better work now than before | 1 = Agree, 0 = Disagree | Positive sub-score 2 |
| 8 | Days are too short | 1 = Agree, 0 = Disagree | Positive sub-score 2 |
| 9 | Enough to get along | 1 = Agree, 0 = Disagree | Positive sub-score 2 |
| 10 | Family is finest in world | 1 = Agree, 0 = Disagree | Positive sub-score 2 |
| 11 | Have definite work to do | 1 = Agree, 0 = Disagree | Positive sub-score 2 |
| 12 | Just as happy as when younger | 1 = Agree, 0 = Disagree | Positive sub-score 2 |
| 13 | More love and affection now | 1 = Agree, 0 = Disagree | Positive sub-score 2 |
| 14 | Most useful period of life | 1 = Agree, 0 = Disagree | Positive sub-score 2 |
| 15 | Satisfied with family treatment | 1 = Agree, 0 = Disagree | Positive sub-score 2 |
| 16 | Satisfied with work i do | 1 = Agree, 0 = Disagree | Positive sub-score 2 |
| 17 | Some use to those around me | 1 = Agree, 0 = Disagree | Positive sub-score 2 |
| 18 | Wish it could go on forever | 1 = Agree, 0 = Disagree | Positive sub-score 2 |
| 19 | Feel about life accomplishments | 3 = Well satisfied, 2 = Reasonably Satisfied, 1 = Dissatisfied | Positive sub-score 3 |
| 20 | Happiness of marriage | 1 = Very unhappy, 2 = Unhappy, 3 = Average, 4 = Happy | Positive sub-score 3 |
| 21 | How often see family or relative | 0 = No family or relatives, 1 = Less than once a year, 2 = About once a month, 3=Once or twice a week, 4= Every day | Positive sub-score 3 |
| 22 | Your life in general | 4 = Very happy, 3 = Moderately happy, 2 = Average, 1 = Unhappy | Positive sub-score 3 |
| 23 | Have had nervous breakdown | 0 = Not Checked, 1 = Checked | Negative sub-score 1 |
| 24 | Worry about health | 0 = Not Checked, 1 = Checked | Negative sub-score 1 |
| 25 | Badly flustered when hurried | 1 = Agree, 0 = Disagree | Negative sub-score 2 |
| 26 | Dreariest time of life | 1 = Agree, 0 = Disagree | Negative sub-score 2 |
| 27 | Family always trying to boss | 1 = Agree, 0 = Disagree | Negative sub-score 2 |
| 28 | Family does not really care | 1 = Agree, 0 = Disagree | Negative sub-score 2 |
| 29 | Happier if could see friends mor | 1 = Agree, 0 = Disagree | Negative sub-score 2 |
| 30 | Haven`t a cent in world | 1 = Agree, 0 = Disagree | Negative sub-score 2 |
| 31 | Just able to make ends meet | 1 = Agree, 0 = Disagree | Negative sub-score 2 |
| 32 | Just no point in living | 1 = Agree, 0 = Disagree | Negative sub-score 2 |
| 33 | Less and less reason to live | 1 = Agree, 0 = Disagree | Negative sub-score 2 |
| 34 | Life could be happier | 1 = Agree, 0 = Disagree | Negative sub-score 2 |
| 35 | Life full of worry | 1 = Agree, 0 = Disagree | Negative sub-score 2 |
| 36 | Life not very useful | 1 = Agree, 0 = Disagree | Negative sub-score 2 |
| 37 | Lonely much of time | 1 = Agree, 0 = Disagree | Negative sub-score 2 |
| 38 | Never dreamed could be as lonely | 1 = Agree, 0 = Disagree | Negative sub-score 2 |
| 39 | No longer do useful work | 1 = Agree, 0 = Disagree | Negative sub-score 2 |
| 40 | No one to talk to | 1 = Agree, 0 = Disagree | Negative sub-score 2 |
| 41 | No work to look forward to | 1 = Agree, 0 = Disagree | Negative sub-score 2 |
| 42 | Wish family paid more attention | 1 = Agree, 0 = Disagree | Negative sub-score 2 |

Summary scores were calculated as follows:

- Positive sub-score 1 was calculated by summing items 1 to 3.
- Positive sub-score 2 was calculated by summing items 4 to 18.
- Positive sub-score 3 was calculated by scaling and then averaging items 19-22.
  Positive total score was calculated by first scaling then sub-score 1, 2 and 3.
- Negative sub-score 1 was calculated by summing items 23-24.
- Negative sub-score 2 was calculated by summing items 25-42.
- Negative total score was calculated by first scaling then sub-score 1 and 2.

Some of the individual items of the questionnaires were not included because they presented too many missing values and others were excluded because they did not correlate with the expected direction of other items of similar valence.
