## Supplemental doc 2 for "Psychosocial experiences are associated with human brain mitochondrial biology"

**BLSA proteomics**

**Tissue Processing**

Each sample was individually homogenized in 300 uL of urea lysis buffer (8M urea, 100 mM NaHPO4, pH 8.5), including 5 uL (100x stock) HALT protease and phosphatase inhibitor cocktail (Pierce) essentially as previously described (1-4). All homogenization was performed using a Bullet Blender (Next Advance) according to manufacturer protocols. Briefly, each tissue piece was added to Urea lysis buffer in a 1.5 mL Rino tube (Next Advance) harboring 750 mg stainless steel beads (0.9-2 mm in diameter) and blended twice for 5 minute intervals in the cold room (4°C). Protein supernatants were transferred to 1.5 mL Eppendorf tubes and sonicated (Sonic Dismembrator, Fisher Scientific) 3 times for 5 s with 15 s intervals of rest at 30% amplitude to disrupt nucleic acids and subsequently vortexed. Protein concentration was determined by the bicinchoninic acid (BCA) method, and samples were frozen in aliquots at −80°C. Protein homogenates (50ug) treated with 1 mM dithiothreitol (DTT) at 25°C for 30 minutes, followed by 5 mM iodoacetimide (IAA) at 25°C for 30 minutes in the dark. Protein mixture (120 ug each) was digested overnight with 1:100 (w/w) lysyl endopeptidase (Wako) at room temperature. The samples were then diluted with 50 mM NH4HCO3 to a final concentration of less than 2M urea and then and further digested overnight with 1:50 (w/w) trypsin (Promega) at 25°C. Resulting peptides were desalted with a Sep-Pak C18 column (Waters) and dried under vacuum.

**Tandem Mass Tag (TMT) Labeling**

Peptides were reconstituted in 100ul of 100mM triethyl ammonium bicarbonate (TEAB) and labeling was performed as previously described (1, 2) using TMTPro isobaric tags (Thermofisher Scientific, A44520 Lot# VH311511 ). Briefly, the TMT labeling reagents were equilibrated to room temperature, and anhydrous ACN (200 μL) was added to each reagent channel. Each channel was gently vortexed for 5 min, and then 20 μL from each TMT channel was transferred to the peptide solutions and allowed to incubate for 1 h at room temperature. The reaction was quenched with 5% (vol/vol) hydroxylamine (5 μl) (Pierce). All 16 channels were then combined and dried by SpeedVac (LabConco) to approximately 100 μL and diluted with 1 mL of 0.1% (vol/vol) TFA, then acidified to a final concentration of 1% (vol/vol) FA and 0.1% (vol/vol) TFA. Peptides were desalted with a 60 mg HLB plate (Waters). The eluates were then dried to completeness.

**High pH Fractionation**

High pH fractionation was performed essentially as described with slight modification (3). Dried samples were re-suspended in high pH loading buffer (0.07% vol/vol NH4OH, 0.045% vol/vol FA, 2% vol/vol ACN) and loaded onto a Water’s BEH (2.1mm x 150 mm with 1.7 µm beads). A Thermo Vanquish UPLC system was used to carry out the fractionation. Solvent A consisted of 0.0175% (vol/vol) NH4OH, 0.01125% (vol/vol) FA, and 2% (vol/vol) ACN; solvent B consisted of 0.0175% (vol/vol) NH4OH, 0.01125% (vol/vol) FA, and 90% (vol/vol) ACN. The sample elution was performed over a 25 min gradient with a flow rate of 0.6 mL/min with a gradient from 0 to 50% B. A total of 96 individual equal volume fractions were collected across the gradient and dried to completeness using a vacuum centrifugation.

**Liquid Chromatography Tandem Mass Spectrometry**

All samples were analyzed on the Evosep One system using an in-house packed 15 cm, 75 μm i.d. capillary column with 1.9 μm Reprosil-Pur C18 beads (Dr. Maisch, Ammerbuch, Germany) using the pre-programmed 21 min gradient (60 samples per day) essentially as described (5). Mass spectrometry was performed with a high-field asymmetric waveform ion mobility spectrometry (FAIMS) Pro equipped Orbitrap Eclipse (Thermo) in positive ion mode using data-dependent acquisition with 2 second top speed cycles. Each cycle consisted of one full MS scan followed by as many MS/MS events that could fit within the given 2 second cycle time limit. MS scans were collected at a resolution of 120,000 (410-1600 m/z range, 4x10^5 AGC, 50 ms maximum ion injection time, FAIMS compensation voltage of -45). All higher energy collision-induced dissociation (HCD) MS/MS spectra were acquired at a resolution of 30,000 (0.7 m/z isolation width, 35% collision energy, 1.25×10^5 AGC target, 54 ms maximum ion time, TurboTMT on). Dynamic exclusion was set to exclude previously sequenced peaks for 20 seconds within a 10-ppm isolation window.

**Database Search and Quantification**

All raw files were searched using Thermo's Proteome Discoverer suite (version 2.4.1.15) with Sequest HT. The spectra were searched against a human uniprot database downloaded August 2020 (86395 target sequences). Search parameters included 10ppm precursor mass window, 0.05 Da product mass window, dynamic modifications methione (+15.995 Da), deamidated asparagine and glutamine (+0.984 Da), phosphorylated serine, threonine and tyrosine (+79.966 Da), and static modifications for carbamidomethyl cysteines (+57.021 Da) and N-terminal and Lysine-tagged TMTPro (+ 304.207 Da). Percolator was used to filter PSMs to 0.1%. Peptides were grouped using strict parsimony and only razor and unique peptides were used for protein level quantitation. Reporter ions were quantified from MS2 scans using an integration tolerance of 20 ppm with the most confident centroid setting. Only unique and razor (i.e., parsimonious) peptides were considered for quantification. Raw data can be accessed at <https://www.synapse.org/#!Synapse:syn39213792>

**Batch correction and data pre-processing**

There were 9,383 total high abundance, master proteins identified across the seven TMT batches. For subsequent analyses, only proteins quantified in >50% of samples were included (n=7,419 proteins). Proteins were normalized as a ratio dividing by the central tendency of pooled standards (Global Internal Standards, GIS) and log2 transformed. Batch correction was completed using an iterative median polish algorithm for removing technical variance across TMT batches, Tunable Approach for Median Polish of Ratio (6). Multidemensional scaling (MDS) plots were used to visualize batch-derived variation before and after TAMPOR batch correction. Outliers were detected and removed based on network connectivity; that is, samples that were greater than 3 standard deviations away from the mean, as described (6). To remove potentially confounding covariates of age, sex and post-mortem interval (PMI), non-parametric bootstrap regression was performed in each region separately. Each trait was subtracted times the median coefficient from 1000 iterations of fitting for each protein, while protecting for diagnosis (control, AsymAD, AD) as previously described (4).

**References**

1.	Johnson, E. C. B., Dammer, E. B., Duong, D. M., Ping, L., Zhou, M., Yin, L., Higginbotham, L. A., Guajardo, A., White, B., Troncoso, J. C., Thambisetty, M., Montine, T. J., Lee, E. B., Trojanowski, J. Q., Beach, T. G., Reiman, E. M., Haroutunian, V., Wang, M., Schadt, E., Zhang, B., Dickson, D. W., Ertekin-Taner, N., Golde, T. E., Petyuk, V. A., De Jager, P. L., Bennett, D. A., Wingo, T. S., Rangaraju, S., Hajjar, I., Shulman, J. M., Lah, J. J., Levey, A. I., and Seyfried, N. T. (2020) Large-scale proteomic analysis of Alzheimer's disease brain and cerebrospinal fluid reveals early changes in energy metabolism associated with microglia and astrocyte activation. *Nat Med* **26**, 769-780

2.	Ping, L., Duong, D. M., Yin, L., Gearing, M., Lah, J. J., Levey, A. I., and Seyfried, N. T. (2018) Global quantitative analysis of the human brain proteome in Alzheimer's and Parkinson's Disease. *Sci Data* **5**, 180036

3.	Ping, L., Kundinger, S. R., Duong, D. M., Yin, L., Gearing, M., Lah, J. J., Levey, A. I., and Seyfried, N. T. (2020) Global quantitative analysis of the human brain proteome and phosphoproteome in Alzheimer's disease. *Sci Data* **7**, 315

4.	Johnson, E. C. B., Carter, E. K., Dammer, E. B., Duong, D. M., Gerasimov, E. S., Liu, Y., Liu, J., Betarbet, R., Ping, L., Yin, L., Serrano, G. E., Beach, T. G., Peng, J., De Jager, P. L., Haroutunian, V., Zhang, B., Gaiteri, C., Bennett, D. A., Gearing, M., Wingo, T. S., Wingo, A. P., Lah, J. J., Levey, A. I., and Seyfried, N. T. (2022) Large-scale deep multi-layer analysis of Alzheimer’s disease brain reveals strong proteomic disease-related changes not observed at the RNA level. *Nature Neuroscience* **25**, 213-225

5.	Bekker-Jensen, D. B., Martínez-Val, A., Steigerwald, S., Rüther, P., Fort, K. L., Arrey, T. N., Harder, A., Makarov, A., and Olsen, J. V. (2020) A Compact Quadrupole-Orbitrap Mass Spectrometer with FAIMS Interface Improves Proteome Coverage in Short LC Gradients. *Mol Cell Proteomics* **19**, 716-729

6.	Johnson, E. C. B., Carter, E. K., Dammer, E. B., Duong, D. M., Gerasimov, E. S., Liu, Y., Liu, J., Betarbet, R., Ping, L., Yin, L., Serrano, G. E., Beach, T. G., Peng, J., De Jager, P. L., Haroutunian, V., Zhang, B., Gaiteri, C., Bennett, D. A., Gearing, M., Wingo, T. S., Wingo, A. P., Lah, J. J., Levey, A. I., and Seyfried, N. T. (2022) Large-scale deep multi-layer analysis of Alzheimer's disease brain reveals strong proteomic disease-related changes not observed at the RNA level. *Nat Neurosci* **25**, 213-225
